# The aggregation potential of Zika virus proteome

**DOI:** 10.1101/2022.03.26.485915

**Authors:** Rajanish Giri, Taniya Bhardwaj, Kumar Udit Saumya, Kundlik Gadhave, Shivani K Kapuganti, Nitin Sharma

## Abstract

The ability of human encoded soluble proteins to convert into amyloid fibrils is now recognized as a generic phenomenon in several human illnesses. Typically, such disease causal proteins/peptides consist of aggregation-prone regions (APR) that make them susceptible to misfolding and assemble into highly ordered β-sheet rich fibrils, distinct from their native soluble state. Here, we show that the zika virus (ZIKV) consists of several such aggregation prone hotspots spread across its entire proteome. Using a combination of high-accuracy prediction tools, we identified APRs in both structural and non-structural proteins of ZIKV. Furthermore, we have experimentally validated the bioinformatic results by subjecting the ZIKV proteins and peptides to artificial aggregation inducing environment. Using a combination of dye-based assays (ThT and ANS) and microscopy techniques (HR-TEM and AFM), we further characterized the morphological features of amyloid-like fibrils. We found that Envelope domain III (EDIII) protein, NS1 *β*-roll peptide, membrane-embedded signal peptide 2K, and cytosolic region of NS4B protein to be highly aggregating in the experimental setup. Our findings also pave the way for an extensive and detailed functional analysis of these predicted APRs in the future to enhance our understanding of the role played by amyloids in the pathogenesis of flavivirus.

**Graphical Abstract:** 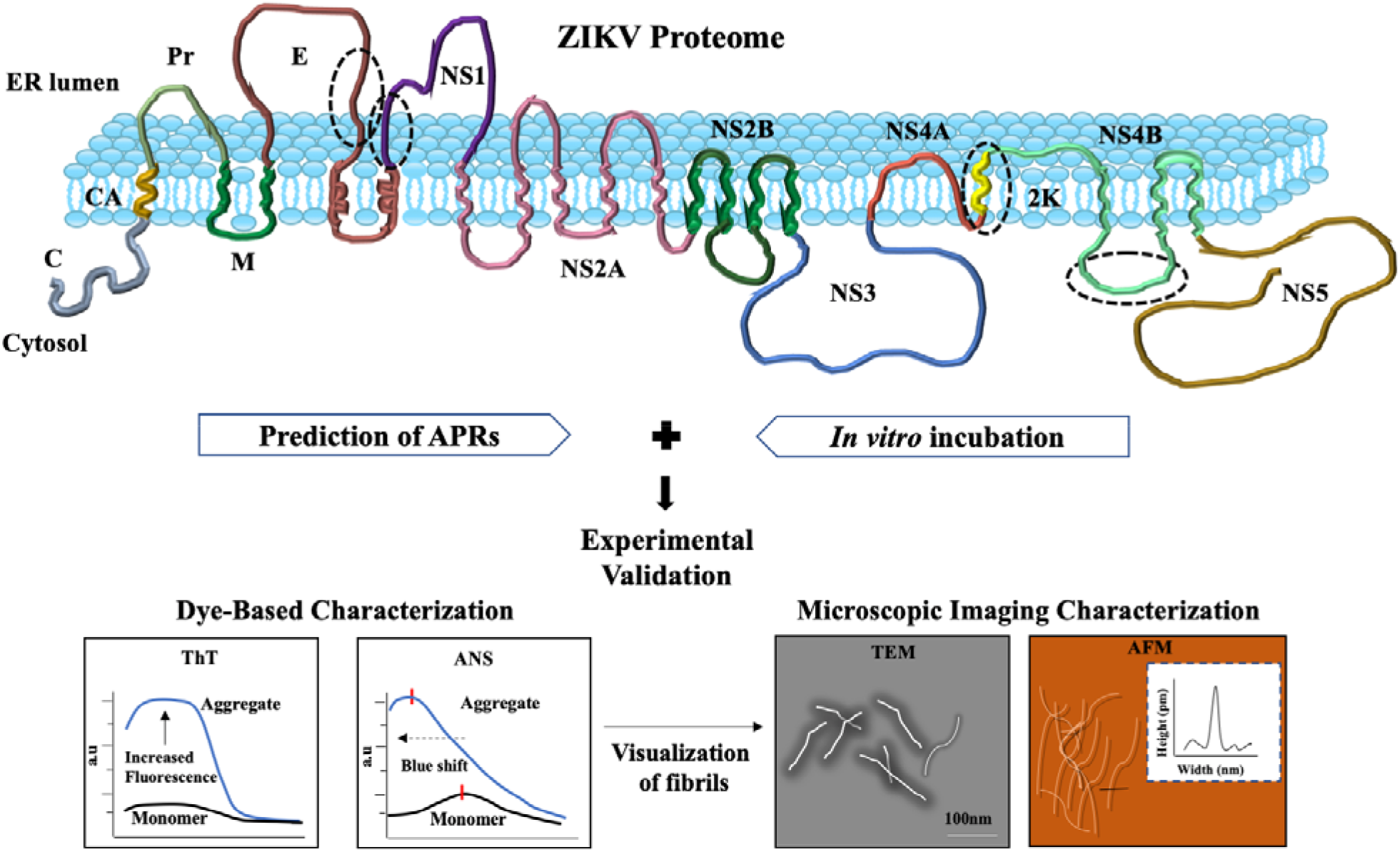

## Introduction

ZIKV is an arthropod-borne RNA virus belonging to the family *flaviviridae*. The onset of ZIKV infection is associated with mild symptoms and rarely requires hospitalization.^1^ However, a significant threat of its infection lies in its neurotropic behavior as a wide range of human cells are highly permissive of its replication. Severe neurological complications such as microcephaly, Guillain-Barré syndrome, and Congenital Zika Syndrome are therefore linked to ZIKV infection.^2^ A few reports have investigated the mechanisms of pathogenesis by ZIKV leading to neurodevelopmental abnormalities. It employs different modes for neuronal cell death, such as disruption of mitochondrial dynamics and its network structure and induction of the release of neurotoxic factors from infected cells to the uninfected ones.^3,4^ In addition, ZIKV causes more rapid neural apoptosis in infants than in older mice models.^5^

Several neurological disorders are attributed to protein misfolding and aggregation into amyloid fibrils, contributing to an extensive array of human diseases that are becoming more prevalent. Alzheimer’s disease (AD), Parkinson’s disease, and Huntington’s disease are some extensively studied proteopathic disorders, primarily caused by the formation and deposition of amyloidogenic proteins.^6^ Typically, such disease causal proteins and peptides undergo misfolding and assemble into highly ordered β-sheet rich fibrils. In this process, aggregation-nucleating regions, also described as aggregation prone regions (APRs) within protein sequence, predispose them to oligomerize into insoluble precursor proto-fibrils. Several of such proto-fibrillar units assemble that eventually mature into amyloid fibrils.^7^ These amyloid fibrils accumulate in tissues and subsequently lead to cellular damage and dysfunction, specific to the kind of protein involved. Viruses are recently known to affect the causal proteins of many neurodegenerative disorders like AD. Viral proteins are also investigated to form amyloids – functional as well as toxic to the host.^8,9^ Thus, an increasing cohort of viral diseases are now linked with amyloidogenic disorders, but surprisingly, this phenomenon in flaviviral infections is yet to be recognized.

Previously, in this context, our group showed the formation of amyloid-like fibrils by ZIKV capsid anchor, which adopts β-sheet conformation, and the aggregates are cytotoxic to mammalian cells.^10^ In that direction, this is important to investigate the aggregation propensity of as many Zika proteins as possible. In this manuscript, we have highlighted multiple hotspots with a high propensity to aggregate across the ZIKV proteome. Our analysis is further validated by aggregation of Envelope domain III (EDIII) protein, NS1 *β*-roll peptide (1-30 residues of NS1 protein), 2K peptide, and NS4B-CR peptide (NS4B cytosolic region – 131-169 residues) through artificially prepared aggregation inducing environment. Our findings here indicate yet another probable mechanism through which ZIKV may impart its pathogenic effect on infected cells.

## Results

Numerous proteopathic disorders are associated with protein misfolding and aggregation.^11–13^ Extensive studies on these pathogenic proteins have led to the recognition of common properties and structural features by virtue of which proteins sustain properties to form aggregates. Thus, APRs of any given polypeptide sequence can now be predicted with high precision tools and software available freely for users. In this study, we have used four different open-source computational tools, namely TANGO, AGGRESCAN, MetAmyl, and FoldAmyloid, to predict the APRs in the ZIKV proteome.^14^ We have also utilized CamSol to identify the hydrophobic regions in ZIKV proteins (**Figures 1** and **2**, and **Tables 1** and **2**).

**Table 1:** Aggregation prone regions in structural proteins of ZIKV. The overlapping regions predicted by different software are highlighted in bold.

| **ZIKV proteins** | **TANGO** | **FoldAmyloid** | **AGGRESCAN** | **MetAmyl** | **CamSol** |
| --- | --- | --- | --- | --- | --- |
| **C**<br>**(104 residues)** | **46-54** | 9-17, **45-58**,<br>64-68, 89-96 | 10-19, 35-41,<br>**44-60**, 62-69,<br>78-83, 89-95 | 11-18, 20-25,<br>**42-52, 54-59** | **47-56** |
| **CA**<br>**(18 residues)** | **6-17** | **7-15** | **6-16** | **4-14** | **7-16** |
| **prM**<br>**(168**<br>**residues)** | **9-14, 72-80,**<br>**134-147,**<br>**154-163** | **9-15, 36-40,**<br>66-70, **74-80,**<br>120-124,<br>127-131,<br>**136-146,**<br>**154-165** | **9-14, 22-31,**<br>**73-80, 120-**<br>**128, 135-148,**<br>**152-164** | 22-41, **69-82,**<br>95-100, 125-130, **135-145,**<br>**149-163** | **68-80, 136-**<br>**145, 154-165** |
| **E**<br>**(500**<br>**residues)** | **197-201,**<br>**250-255,**<br>**381-388,**<br>**458-473,**<br>**480-498** | 1-5, 21-25,<br>136-142,<br>**196-201,**<br>211-217,<br>**250-254,**<br>279-283,<br>**382-386,**<br>442-447,<br>**457-472,**<br>**481-486,**<br>**489-496** | 15-30, 43-51,<br>108-121, 137-144, 177-181,<br>**194-200, 249-**<br>**257, 298-313,**<br>321-327, 339-348, 351-358,<br>**379-389, 428-**<br>**475, 480-500** | 1-8, 18-26,<br>28-36, 40-53,<br>56-61, 86-95,<br>109-126,<br>138-147,<br>160-166,<br>**195-200,**<br>237-242,<br>**247-264,**<br>296-314,<br>319-331,<br>346-356,<br>**379-390,**<br>409-414,<br>427-438,<br>**459-471,**<br>**487-498** | 30-34, **197-**<br>**201, 301-311,**<br>320-327,<br>**381-386,**<br>443-447,<br>**458-472,**<br>**481-496** |

**Table 2:**
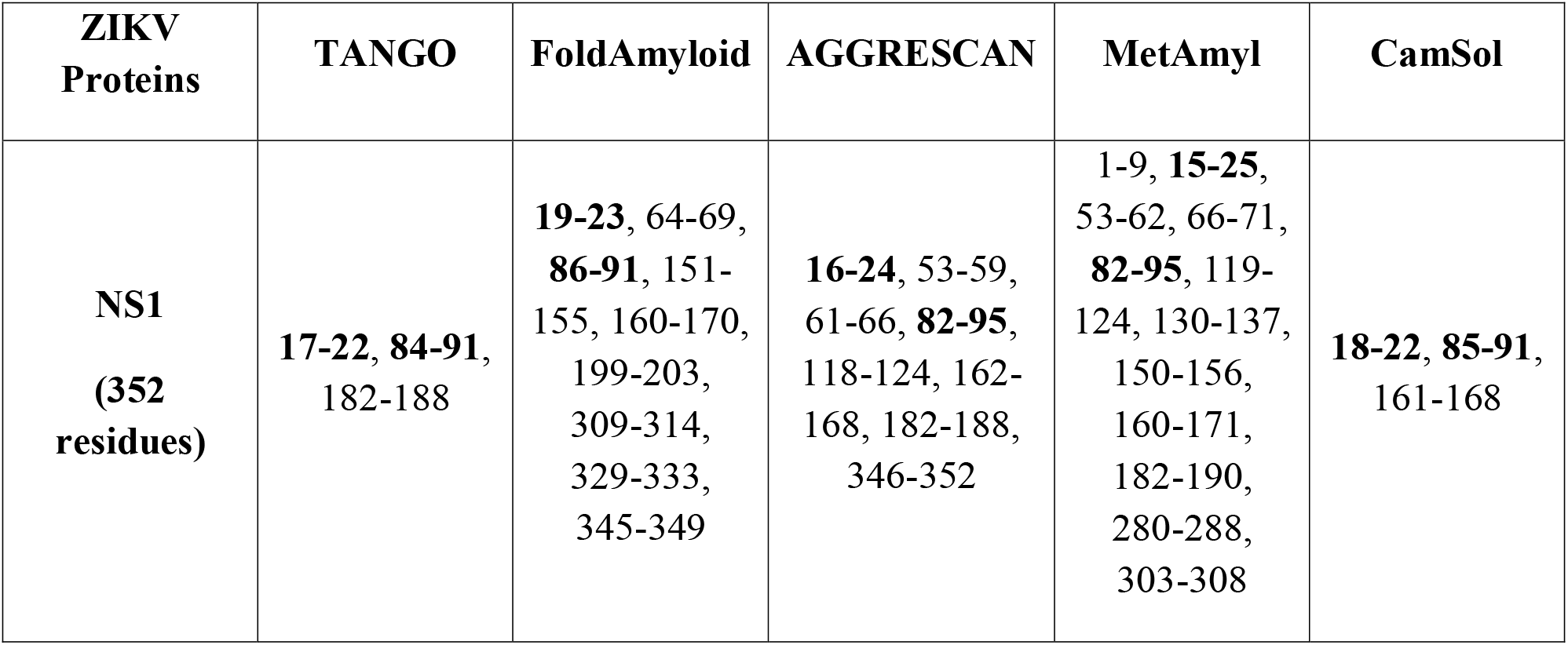
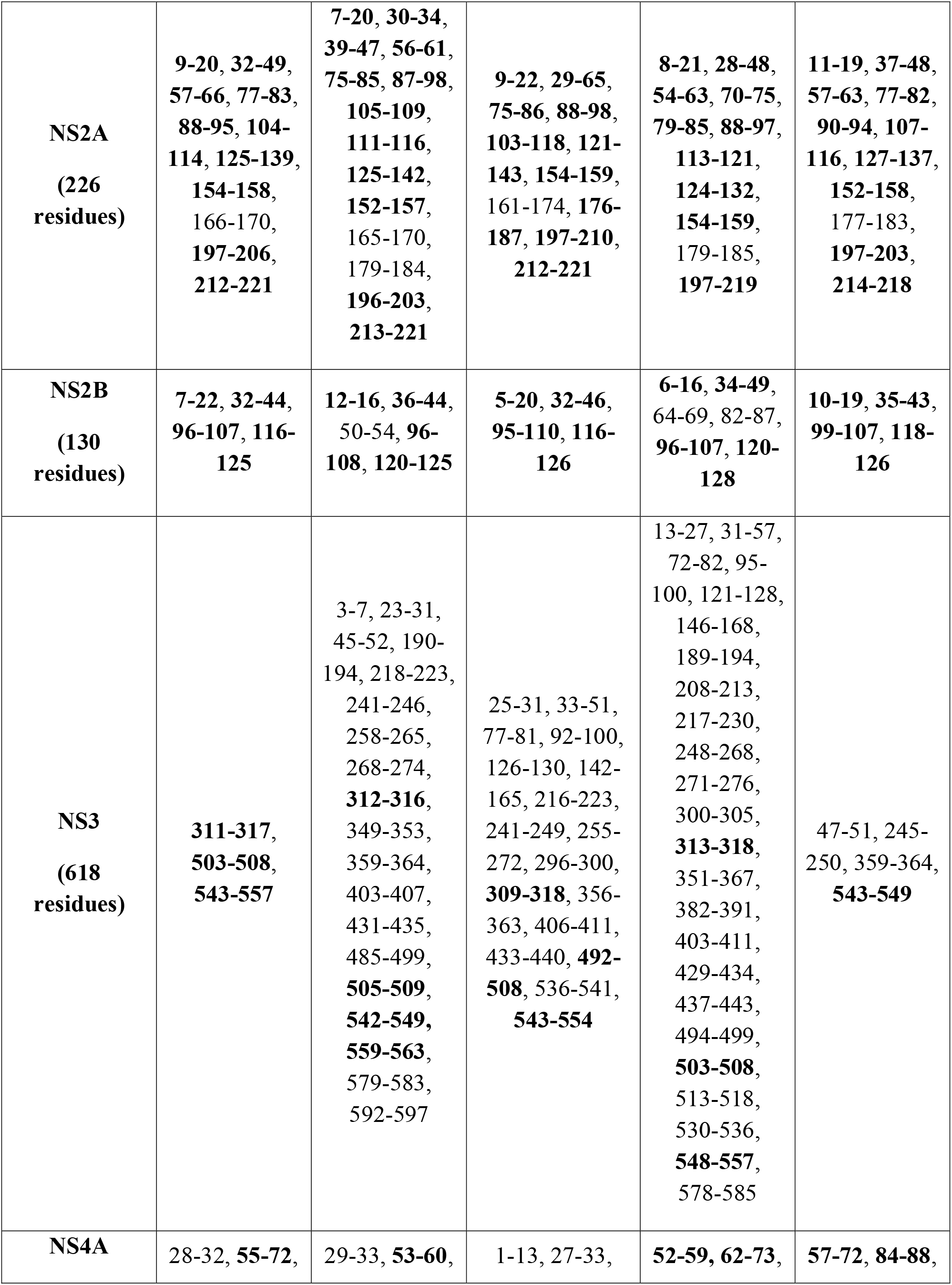
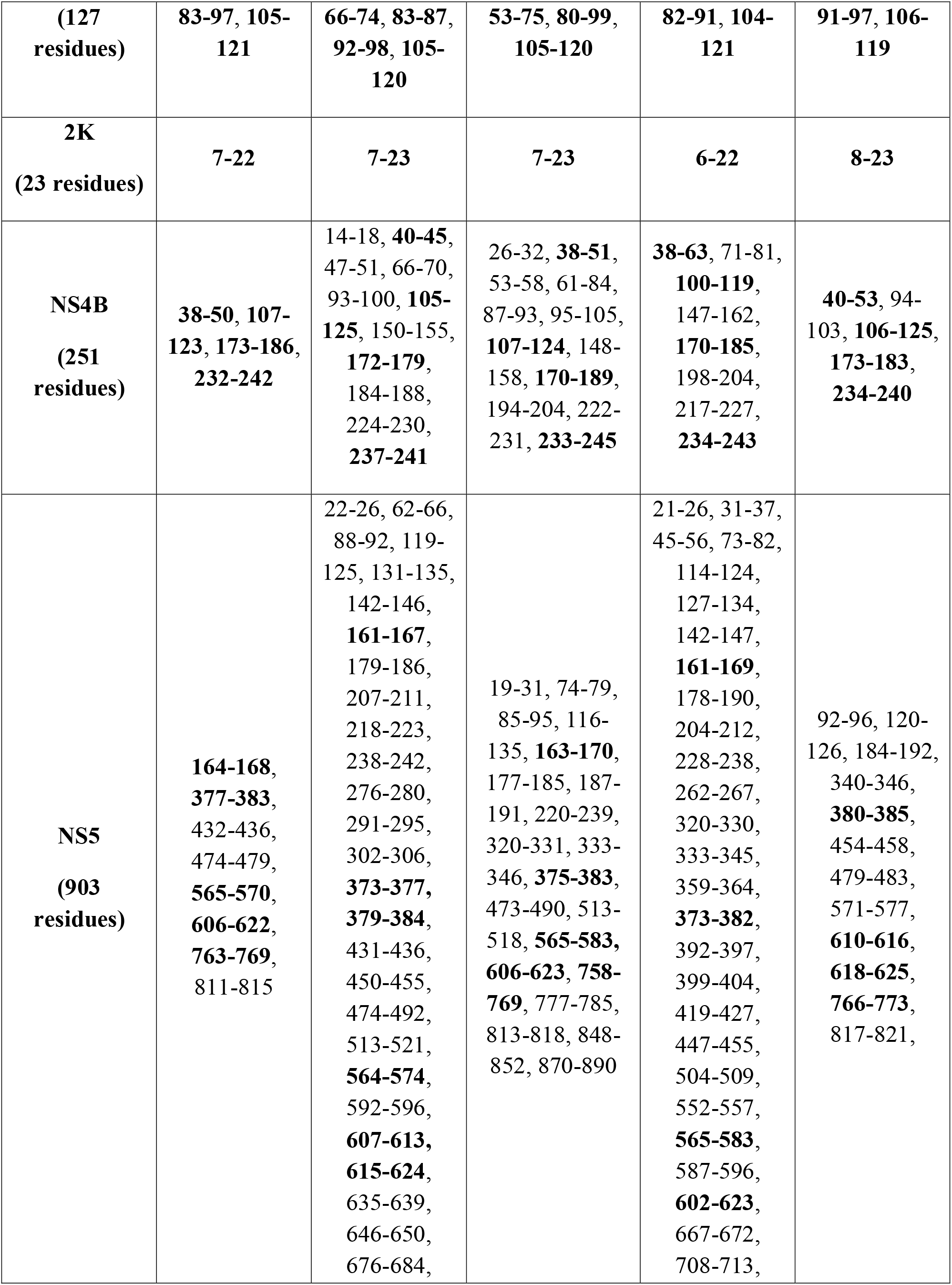
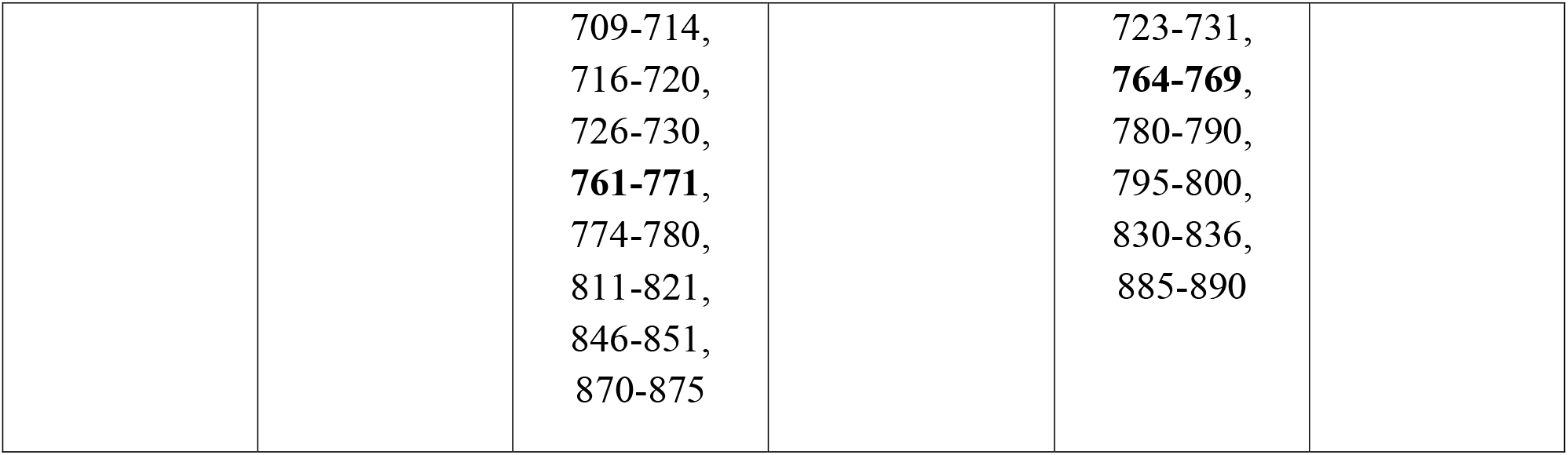
Aggregation prone regions in ZIKV non-structural proteins. The overlapping regions are highlighted in bold.

| **ZIKV Proteins** | **TANGO** | **FoldAmyloid** | **AGGRESCAN** | **MetAmyl** | **CamSol** |
| --- | --- | --- | --- | --- | --- |
| **NS1**<br>**(352 residues)** | **17-22, 84-91,**<br>182-188 | **19-23,** 64-69,<br>**86-91,** 151-155, 160-170,<br>199-203,<br>309-314,<br>329-333,<br>345-349 | **16-24,** 53-59,<br>61-66, **82-95,**<br>118-124, 162-168, 182-188,<br>346-352 | 1-9, **15-25,**<br>53-62, 66-71,<br>**82-95,** 119-124, 130-137,<br>150-156,<br>160-171,<br>182-190,<br>280-288,<br>303-308 | **18-22, 85-91,**<br>161-168 |
| **NS2A**<br>**(226 residues)** | **9-20, 32-49, 57-66, 77-83, 88-95, 104-114, 125-139, 154-158, 166-170, 197-206, 212-221** | **7-20, 30-34, 39-47, 56-61, 75-85, 87-98, 105-109, 111-116, 125-142, 152-157, 165-170, 179-184, 196-203, 213-221** | **9-22, 29-65, 75-86, 88-98, 103-118, 121-143, 154-159, 161-174, 176-187, 197-210, 212-221** | **8-21, 28-48, 54-63, 70-75, 79-85, 88-97, 113-121, 124-132, 154-159, 179-185, 197-219** | **11-19, 37-48, 57-63, 77-82, 90-94, 107-116, 127-137, 152-158, 177-183, 197-203, 214-218** |
| **NS2B**<br>**(130 residues)** | **7-22, 32-44, 96-107, 116-125** | **12-16, 36-44, 50-54, 96-108, 120-125** | **5-20, 32-46, 95-110, 116-126** | **6-16, 34-49, 64-69, 82-87, 96-107, 120-128** | **10-19, 35-43, 99-107, 118-126** |
| **NS3**<br>**(618 residues)** | **311-317, 503-508, 543-557** | **3-7, 23-31, 45-52, 190-194, 218-223, 241-246, 258-265, 268-274, 312-316, 349-353, 359-364, 403-407, 431-435, 485-499, 505-509, 542-549, 559-563, 579-583, 592-597** | **25-31, 33-51, 77-81, 92-100, 126-130, 142-165, 216-223, 241-249, 255-272, 296-300, 309-318, 356-363, 406-411, 433-440, 492-508, 536-541, 543-554** | **13-27, 31-57, 72-82, 95-100, 121-128, 146-168, 189-194, 208-213, 217-230, 248-268, 271-276, 300-305, 313-318, 351-367, 382-391, 403-411, 429-434, 437-443, 494-499, 503-508, 513-518, 530-536, 548-557, 578-585** | **47-51, 245-250, 359-364, 543-549** |
| **NS4A** | **28-32, 55-72,** | **29-33, 53-60,** | **1-13, 27-33,** | **52-59, 62-73,** | **57-72, 84-88,** |
| **(127**<br>**residues)** | **83-97, 105-**<br>**121** | **66-74, 83-87,**<br>**92-98, 105-**<br>**120** | **53-75, 80-99,**<br>**105-120** | **82-91, 104-**<br>**121** | **91-97, 106-**<br>**119** |
| **2K**<br>**(23 residues)** | **7-22** | **7-23** | **7-23** | **6-22** | **8-23** |
| **NS4B**<br>**(251**<br>**residues)** | **38-50, 107-**<br>**123, 173-186,**<br>**232-242** | 14-18, **40-45**,<br>47-51, 66-70,<br>93-100, **105-**<br>**125**, 150-155,<br>**172-179**,<br>184-188,<br>224-230,<br>**237-241** | 26-32, **38-51**,<br>53-58, 61-84,<br>87-93, 95-105,<br>**107-124**, 148-158, **170-189**,<br>194-204, 222-231, **233-245** | **38-63**, 71-81,<br>**100-119**,<br>147-162,<br>**170-185**,<br>198-204,<br>217-227,<br>**234-243** | **40-53**, 94-103, **106-125**,<br>**173-183**,<br>**234-240** |
| **NS5**<br>**(903**<br>**residues)** | **164-168,**<br>**377-383,**<br>**432-436,**<br>**474-479,**<br>**565-570,**<br>**606-622,**<br>**763-769,**<br>**811-815** | 22-26, 62-66,<br>88-92, 119-125, 131-135,<br>142-146,<br>**161-167**,<br>179-186,<br>207-211,<br>218-223,<br>238-242,<br>276-280,<br>291-295,<br>302-306,<br>**373-377**,<br>**379-384**,<br>431-436,<br>450-455,<br>474-492,<br>513-521,<br>**564-574**,<br>592-596,<br>**607-613**,<br>**615-624**,<br>635-639,<br>646-650,<br>676-684, | 19-31, 74-79,<br>85-95, 116-135, **163-170**,<br>177-185, 187-191, 220-239,<br>320-331, 333-346, **375-383**,<br>473-490, 513-518, **565-583**,<br>**606-623**, **758-**<br>**769**, 777-785,<br>813-818, 848-852, 870-890 | 21-26, 31-37,<br>45-56, 73-82,<br>114-124,<br>127-134,<br>142-147,<br>**161-169**,<br>178-190,<br>204-212,<br>228-238,<br>262-267,<br>320-330,<br>333-345,<br>359-364,<br>**373-382**,<br>392-397,<br>399-404,<br>419-427,<br>447-455,<br>504-509,<br>552-557,<br>**565-583**,<br>587-596,<br>**602-623**,<br>667-672,<br>708-713, | 92-96, 120-126, 184-192,<br>340-346,<br>**380-385**,<br>454-458,<br>479-483,<br>571-577,<br>**610-616**,<br>**618-625**,<br>**766-773**,<br>817-821, |
|  |  | 709-714,<br>716-720,<br>726-730,<br>**761-771**,<br>774-780,<br>811-821,<br>846-851,<br>870-875 |  | 723-731,<br>**764-769**,<br>780-790,<br>795-800,<br>830-836,<br>885-890 |  |

**Figure 1:**
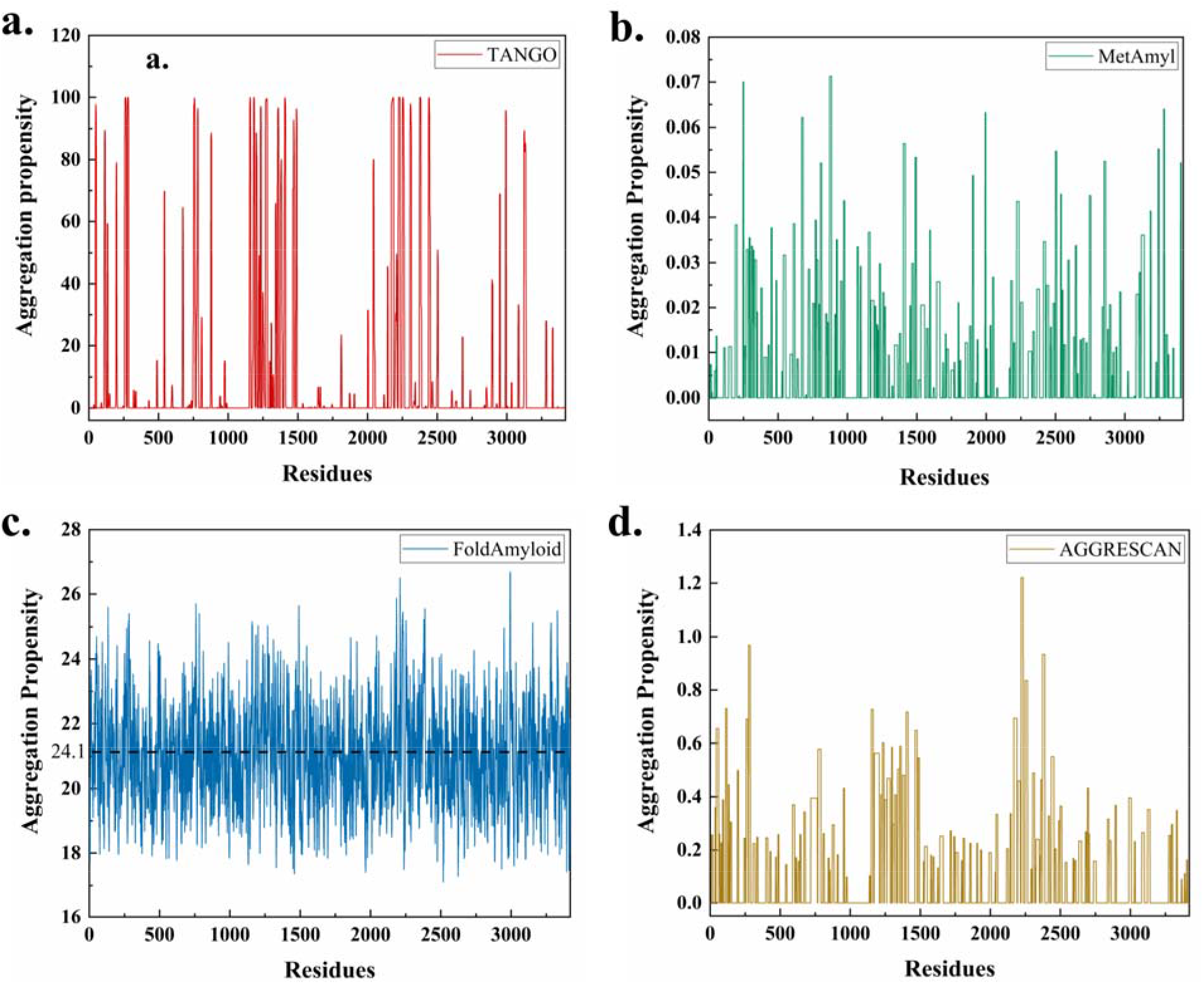
Propensity of aggregation predicted using highly reliable servers: **(A)** TANGO, **(B)** MetAmyl, **(C)** FoldAmyloid, and **(D)** AGGRESCAN in residues of ZIKV proteome.

**Figure 2:**
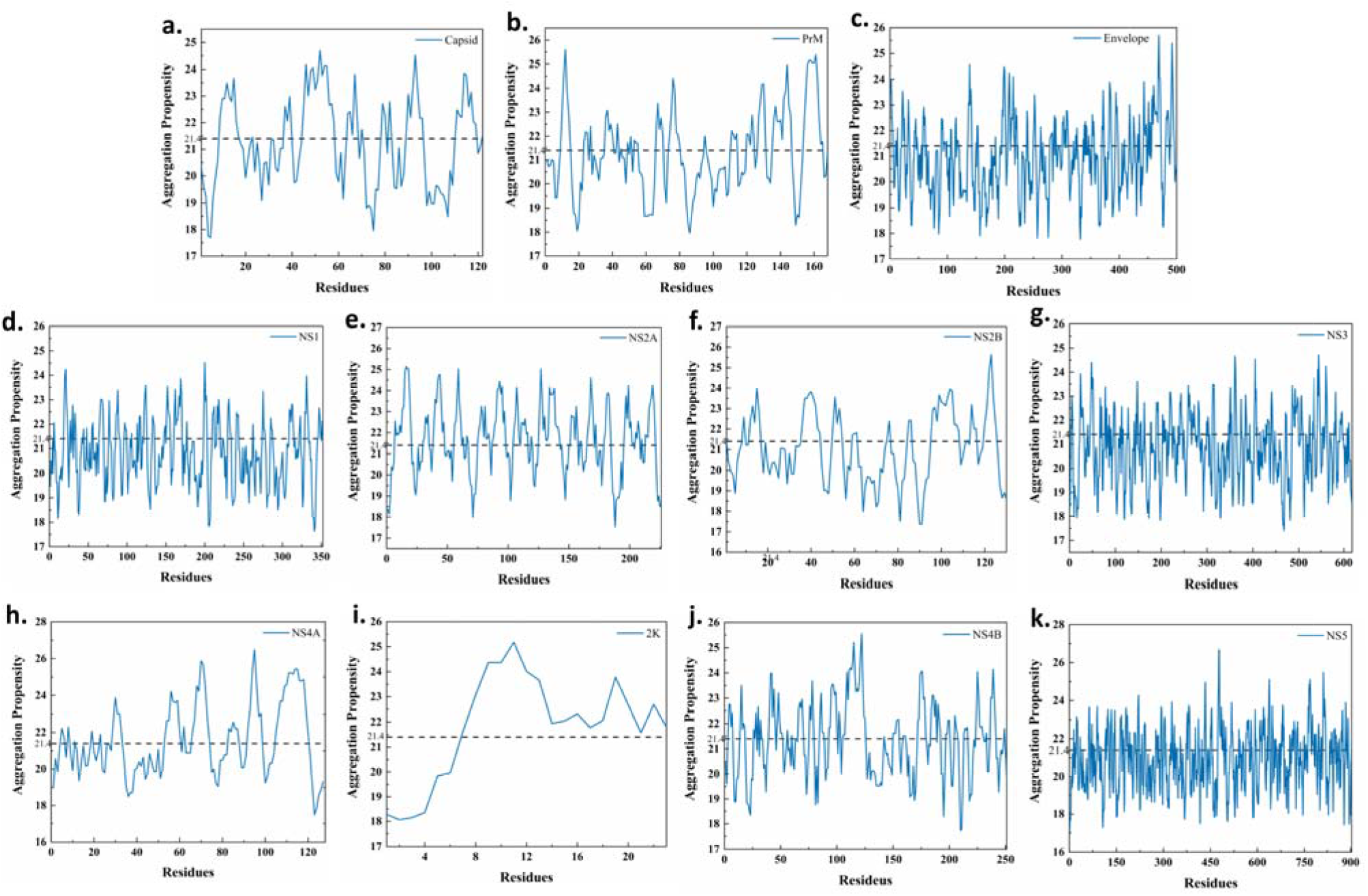
Aggregation prone regions (APRs) predicted by FoldAmyloid in ZIKV (a-c) structural proteins - Capsid, PrM, and Envelope, (i) Corresponds to APRs in 2K peptide, and (d-k) non-structural proteins - NS1, NS2A, NS2B, NS3, NS4A, NS4B, and NS5.

### High aggregation propensity of ZIKV structural proteins

The three structural proteins: Capsid (C), membrane precursor (prM), and Envelope (E), are essential for virus particle formation. Aggregation prone regions in them are predicted under physiological conditions.

The ~12-kDa C protein is synthesized first, and its primary role is to facilitate the packaging of vg-RNA. The 104 amino acid long ZIKV C protein structurally comprises of an intrinsically disordered N-terminal region, four α-helices (α1-α4), and short interspersed loops connecting the helices.^15^ Residues 11-17 are predicted as APR by three prediction tools in our study (**Figure 2a** and **Table 1**). Further, our computational screening also revealed several APRs containing residues 45-52. This particular region comprises of helices α2 and α3, which play a crucial role in stabilizing capsid homodimers through extensive hydrogen-bonding and hydrophobic interactions with surrounding protomer residues.^15^ Also, nearby residues 42 and 43 are reported to be conserved and functionally indispensable for nuclear localization of the capsid protein.^17^ At its C-terminus, C protein is connected to an 18-residue peptide known as capsid anchor (CA). Previously, we have reported the aggregation potential of ZIKV CA and have shown that the amyloid-like aggregates of CA are toxic to mammalian cells.^10^

The structural proteins positioned next to the capsid towards ER lumen are prM and E proteins. The prM protein is composed of pr peptide and M protein, while E protein consists of three domains: EDI, EDII, and EDIII, along with stem and transmembrane regions.^19^ The prM protein plays a crucial role in the transition of non-infectious immature viral particles to infectious mature virion, while E protein serves as the major membrane fusion and host surface binding protein for the virus.^20,21^ During maturation, furin protease assisted cleavage separates pr from M, leaving behind M protein as membrane-embedded while pr fragment falls off.^22^ Our APR scanning in the 168 amino acid long prM protein led us to identify multiple hotspots, especially within residues 74-80, 136-145, and 154-163, which contains high scoring common APRs predicted by all five prediction tools. The 500 amino acid long E protein, which is the largest among all structural proteins, also contains many common APRs such as residues 197-200, 250-254, 381-386, 465-471, and 489-496 (**Figure 2c** and **Table 1**).

### Aggregation prone regions in ZIKV non-structural proteins

Using the same four computational tools, the aggregation prone regions in the seven non-structural proteins and a transmembrane peptide 2K are predicted under physiological conditions.

The first non-structural protein, NS1, is considered essential in host immune surveillance and interaction with host membranes.^23^ It is composed of three domains: N-terminal β-roll (residues 1–30), wing domain (residues 31–180), and C-terminal β-ladder (residues 181–352) in addition to a flexible intertwined loop (residues 91–130) between the β5- and β6-strands of the wing domain.^24^ Our analysis revealed the presence of APRs at its N-terminal and central region. The prediction tools commonly identified the following regions as APRs: residues 16-22 and 86-91; however residues 182-188 are predicted by three servers TANGO, AGGRESCAN, and MetAmyl (**Figure 2d** and **Table 2**). Further, the succeeding proteins in the polyprotein NS2A and NS2B have several common APR hotspots distributed throughout the protein. Flavivirus NS2A protein is a membrane-associated hydrophobic protein engaged in multiple functions besides its involvement in RNA replication.^25^ The NS2B is another membrane-bound protein engaged in interaction with NS3 to form an active NS2B–NS3 protease complex.^26^ In our analysis, we found that the residues 7-20, 29-65 and 75-98, 104-160, and 197-220 consisted of many APRs in NS2A protein while residues 7-16, 32-44, 96-107, and 120-125 forms common APRs in NS2B protein (**Table 2**).

NS3, a bifunctional enzyme, consists of the N-terminal protease domain (residues 1–167) and the C-terminal helicase domain (residues 168–618), essential for polyprotein processing and viral replication, respectively.^27^ According to our observations, MetAmyl has predicted a maximum number of APRs in this protein, while TANGO has predicted the least. Most of the APRs are located in the helicase domain of NS3 protein (Refer to **Table 2** for all APRs in NS3 protein). NS4A and NS4B proteins functions in rearranging the host ER membrane contributing to the formation of virus-induced membranous vesicles.^28,29^ Recently, ZIKV NS4A has been established as a cofactor of NS3 helicase ATPase activity.^30^ From our analysis, we observed that APR distribution was majorly concentrated between the residues 53-73 and 104-121. In ZIKV NS4B protein, residues 38-50, 100-120, and 230-243 contain most of the APRs. In contrast, the 2K transmembrane region of 23 amino acids preceding the NS4B protein is predicted to be highly aggregation prone, having the potential to aggregate in most of its residues.

The largest non-structural protein, NS5, is a complex protein with dual catalytic activity. Its N-terminus (residues 1–252) is characterized by methyl-transferase (MTase) activity, whereas its C-terminal domain (residues 274–892) has the RNA-dependent RNA polymerase (RdRp) activity.^31^ Our analysis has revealed the presence of several hotspots distributed in both these functional domains; however, major APRs are present in the RdRp domain. Several APRs 164-168, 377-382, 565-570, 606-622, and 763-769 residues are located in NS5 protein (see **Table 2** for all APRs).

### 20S Proteasome cleavage sites in ZIKV proteome

We have mapped the 20S proteasome cleavage sites in all the proteins of ZIKV using a SVM-based method known as Pcleavage and a proteasome site predictor NetChop.^32,33^ According to the results, all the ZIKV proteins contain the 20S cleavage sites; however, NetChop has predicted more cleavage sites than the Pcleavage method (**Table S1**).

### In-vitro aggregation of ZIKV proteins and peptides

We experimentally investigated the aggregation behavior of certain proteins and peptides of ZIKV. For this purpose, we have used ThT dye- and ANS dye-based assays to observe the fibrillation process and HR-TEM and AFM to visualize fibril morphology.

#### EDIII protein

The EDIII corresponds to 302-409 residues of full-length E protein. It forms the antigenic determinant residues for host immune responses and, therefore, is considered a potential immunogen in subunit vaccine development.^34^ According to our analysis, it contains potential amyloid-forming segments throughout the region (**Figure 2c** and **Table 1**). TANGO and FoldAmyloid have predicted overlapping APRs in residues 381-388 and 382-386, respectively.

MetAmyl, a meta-predictor, has predicted a different fragment having aggregation propensity from residues 319-331 and residues 379-390 in the EDIII protein region. These observations revealed that the EDIII domain has aggregation propensity, and the regions identified in this study may have an essential role in its amyloid-like fibril formation. Further, in in-vitro experiments, the aggregates formed by the EDIII domain are ThT and ANS positive. We observe a four-fold rise in ThT fluorescence on binding with the mature aggregates at 484 hrs as compared with the EDIII monomer (**figure 3a**). ANS assay also confirms the exposure of hydrophobic residues in the EDIII domain (**figure 3b**) as the incubated sample exhibits an increase in ANS fluorescence and a characteristic blue shift of 23 nm from 536 nm (for EDIII monomer) to 513 nm (484 hr aggregate). The HR-TEM employed to study the morphological features of EDIII aggregates represents distinctive amyloid-like fibrils (**figure 3c**).

**Figure 3:**
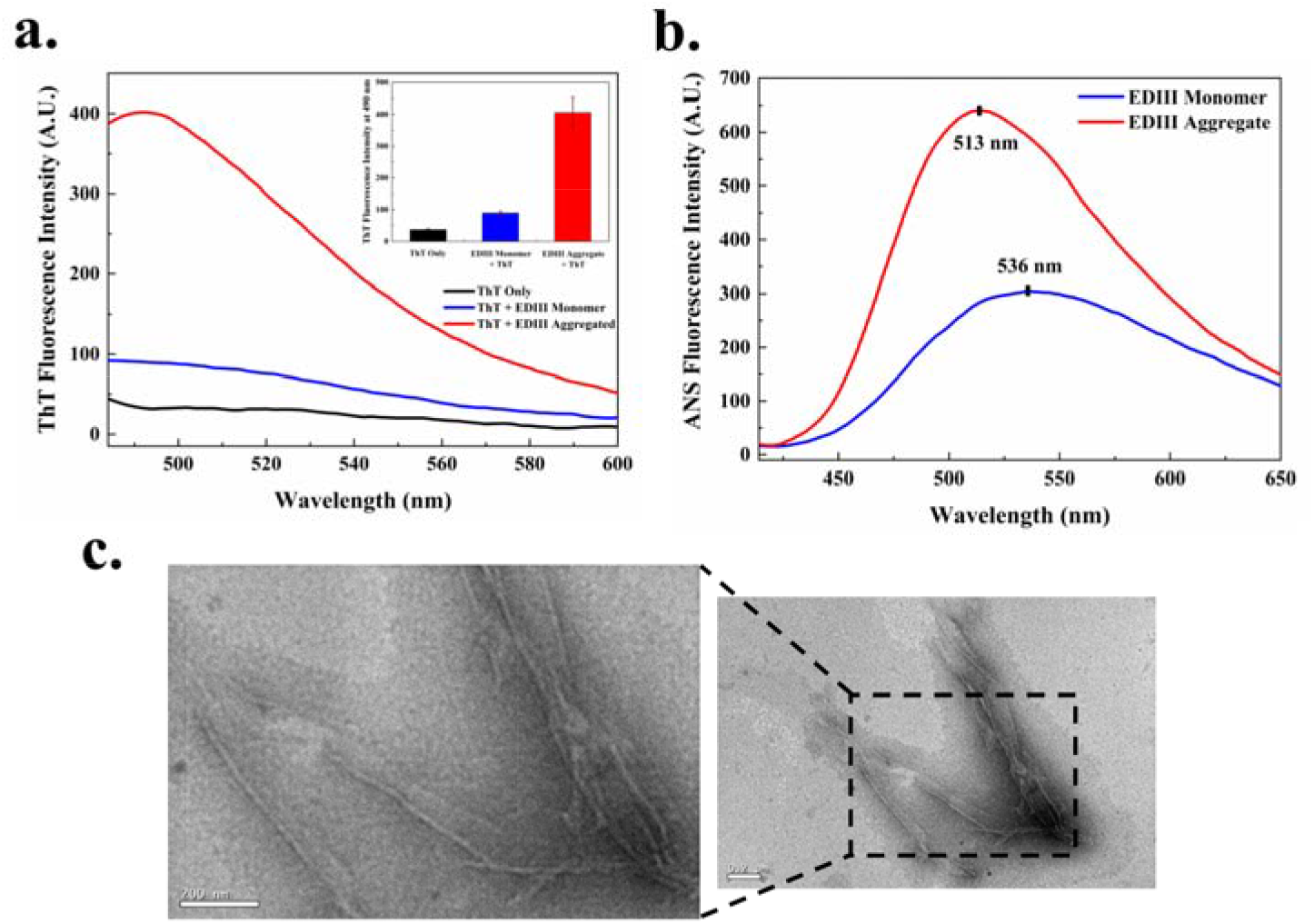
Formation of amyloids-like aggregates by ZIKV EDIII protein. **(a)** ThT fluorescence spectra of EDIII. ~Four-fold increase in ThT fluorescence at 490 nm indicates the formation of amyloid fibrils (484 hrs aggregate). **(b)** ANS fluorescence spectra reveal a blue shift of 26 nm from 536 nm (monomer) to 513 nm (484 hrs aggregate). **(c)** HR-TEM image after 484 hrs shows the amyloid-like fibrillar morphology of aggregated EDIII. The Scale bars represent 200 nm.

#### NS1 *β*-roll peptide

The *β*-roll domain (residues 1-30) of flavivirus NS1 forms a *β*-sheet structure necessary for the protein’s oligomerization and interaction with cellular membranes.^24^ According to the observations, both ThT and ANS dyes bind to the mature fibrils of NS1 *β*-roll peptide. With ThT dye, a ten-fold increase in its fluorescence with the aggregated sample is detected after 24 hrs of incubation (**figure 4a**). It undergoes nucleation-dependent kinetics of amyloidogenesis, studied using ThT dye in this study. **Figure 4b** shows the kinetics of aggregation reaction with a speedy growth phase of ~7 hrs with only a short lag phase of ~1.5 hrs. The halftime (*t*_*50*_) of kinetics reaction is calculated to be 3.89±0.08 hrs. Further, upon binding to hydrophobic regions exposed during aggregate formation, ANS fluorescence shows a blue shift after 24 hrs of incubation. The dye in presence of monomer exhibits an emission maxima at 539.4 nm, which shifts to 495.5 nm on binding to incubated sample (**figure 4c**). The morphological features of mature NS1 *β*-roll amyloid-like fibrils are investigated using both AFM and HR-TEM (**figure 4d–e**). The mature aggregates of 24 hrs appear short fibrillar under HR-TEM. In AFM images obtained after 192 hrs incubation, aggregated fibrils appear long and tangled.

**Figure 4:**
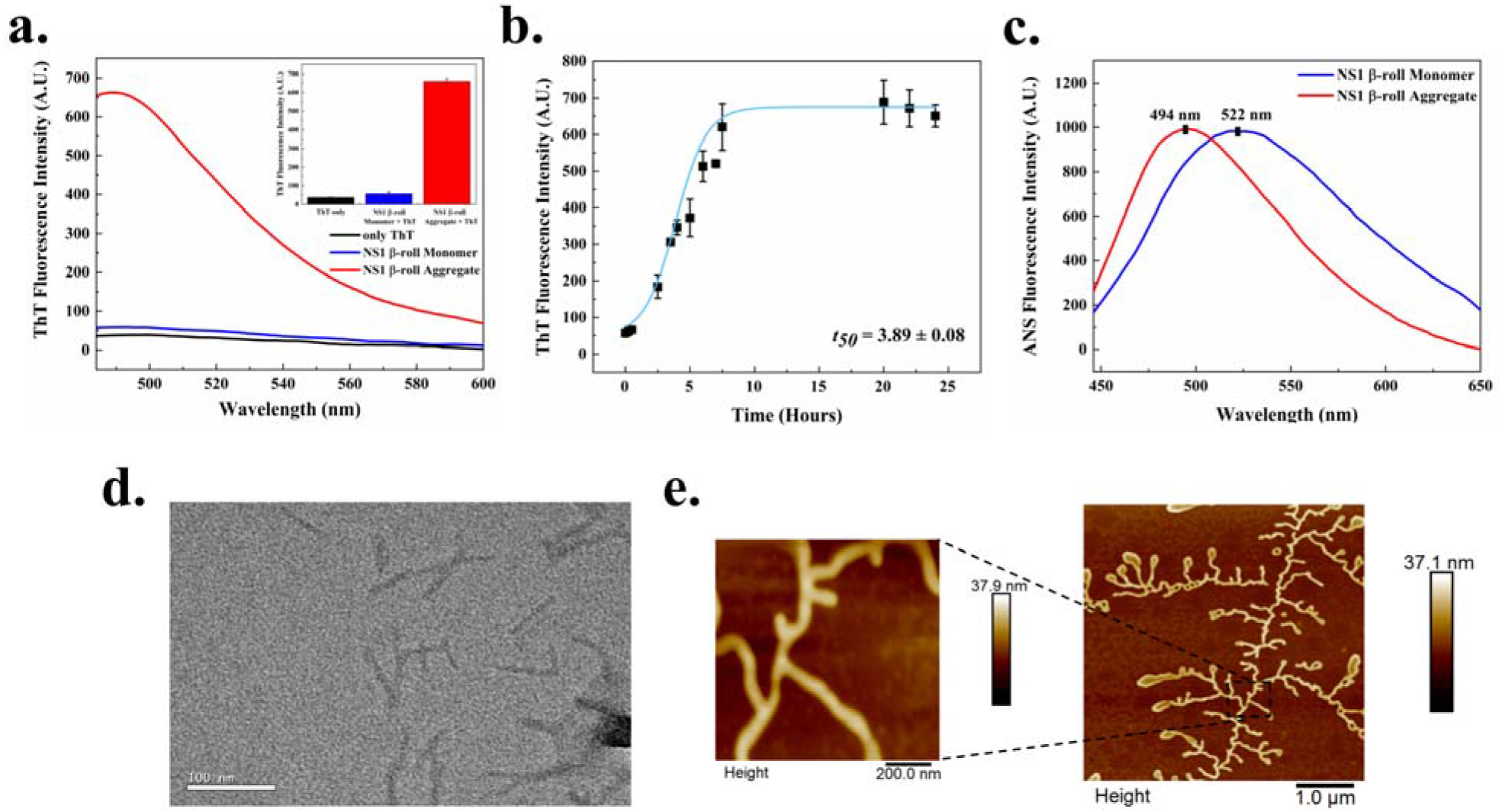
ZIKV NS1 β-roll domain aggregation in-vitro. **(a)** ThT fluorescence scan of NS1 β-roll domain monomer (0 hr incubated) and aggregates (24 hrs aggregates) indicates a ~ten-fold increase in ThT fluorescence at 490 nm **(b)** ThT fluorescence kinetics shows a *t_50_* of 3.89±0.08 hrs for aggregation of NS1 β-roll domain. **(c)** ANS fluorescence produces a blue-shift from 522.3 nm (monomer) to 494.4 nm (aggregate) after 24 hrs. **(d)** HR-TEM image of 24 hrs aggregates. **(e)** AFM image showing the morphology of fibrillar amyloids after 192 hrs aggregates.

#### 2K peptide

The flavivirus 2K peptide is a 23 amino acid fragment between NS4A and NS4B proteins.^35^ ZIKV 2K is predicted to contain a fragment of 7-22 residues as an APR by all the servers used in this study (**Figure 2i,** and **Table 2**). **Figure 5a** depicts the ThT assay wherein the aggregated sample of 2K peptide displays a ~six-fold increase of ThT fluorescence intensity compared to the monomer. With ANS, ~2.5-fold increase in fluorescence intensity of aggregated sample (720 hrs) is observed in comparison with the monomer. The increasing fluorescence was also accompanied by a characteristic blue shift in the emission spectra from 535 nm to 490 nm, further indicating the formation of amyloid fibrils by 2K (**figure 5b**). Furthermore, the HR-TEM images of the 720 hrs incubated sample reveal a typical amyloid structure with dense thread-like long and short interconnected fibrils (**figure 5c**). AFM images shown in **figure 5d** also uncovered the similar fibrillar structures of amyloid aggregates of 2K peptide. **Figure 5e** represents the height profile which quantitatively measures the height and width of the aggregates. These aggregated 2K peptide microscopy images are analogous to previously reported amyloids produced by many cytosolic and transmembrane protein aggregates.

**Figure 5:**
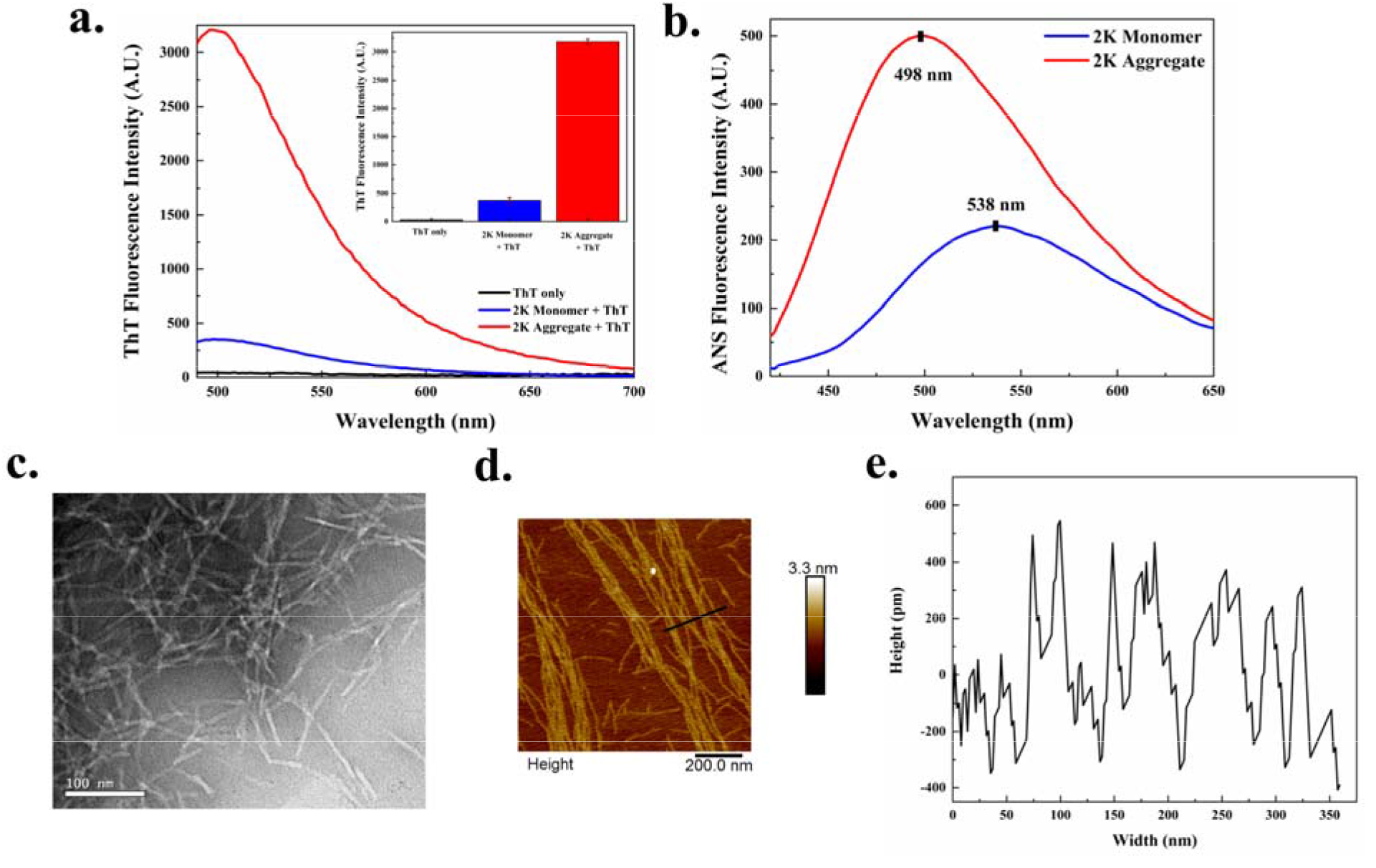
In-vitro aggregation of ZIKV 2K peptide. **(a)** ThT fluorescence spectra of 2K monomer (0 hr incubated) and aggregates show a ~ten-fold increase in ThT fluorescence intensity at 490 nm after 720 hrs. **Inset:** Bar graph represents the ThT fluorescence emission at 490 nm emission wavelength. **(b)** ANS fluorescence emission spectra reveal a characteristic blue-shift from 538 nm (monomer) to 498 nm (720 hrs aggregate). **(c)** HR-TEM and **(d)** AFM images of 2K amyloid fibrils (720 hrs aggregate). The scale bars represent 100 nm and 200 nm for HR-TEM and AFM images. **(e)** The height profile of 2K aggregates is shown in section **(d)**.

#### NS4B-CR peptide

NS4B has a cytosolic region of ~35 amino acids present between the transmembrane domains 3 and 4, which plays a crucial role in replicating the viral genome.^36^ The bioinformatic analysis shows the presence of multiple APRs throughout the NS4B protein, one of which is a five residue APR segment predicted by FoldAmyloid server from 150-155 amino acids spanning the cytosolic region (**Figure 2j** and **Table 2**). MetAmyl also predicted residues 147-162 in NS4B protein to have significant susceptibility to form amyloid aggregates. The amyloid nature of NS4B-CR aggregates is confirmed from **figure 6a,** where a ~three-fold increase in ThT fluorescence intensity is observed at 81 hrs. However, freshly solubilized sample and ThT control have displayed almost parallel ThT fluorescence with no significant difference. Further, its aggregation kinetics depicts a lag phase of about ~17 hrs leading to a ~39 hrs long elongation, eventually phase attaining saturation (**figure 6b**). *t_50_* of kinetics is estimated to be 31.2±0.35 hrs for NS4B-CR aggregation kinetics. Likewise, ANS fluorescence emission is found to be blue-shifted by 7 nm (**figure 6c**). Monomerized NS4B-CR peptide with ANS dye produces a fluorescence emission maxima at 535 nm, while 120 hrs aggregated sample yields a fluorescence emission maxima at 528 nm. Under tapping-mode AFM, various connected fibrils are found in 96 hrs incubated sample (**figure 6e**). The 3-dimensional analysis of the AFM image shown in **figure 6f** identifies the height distribution of fibrils peaking at ~6 nm. Additionally, TEM images of 96 hrs incubated samples uncovered interconnected and overlapping amyloid fibrils of varying diameters.

**Figure 6:**
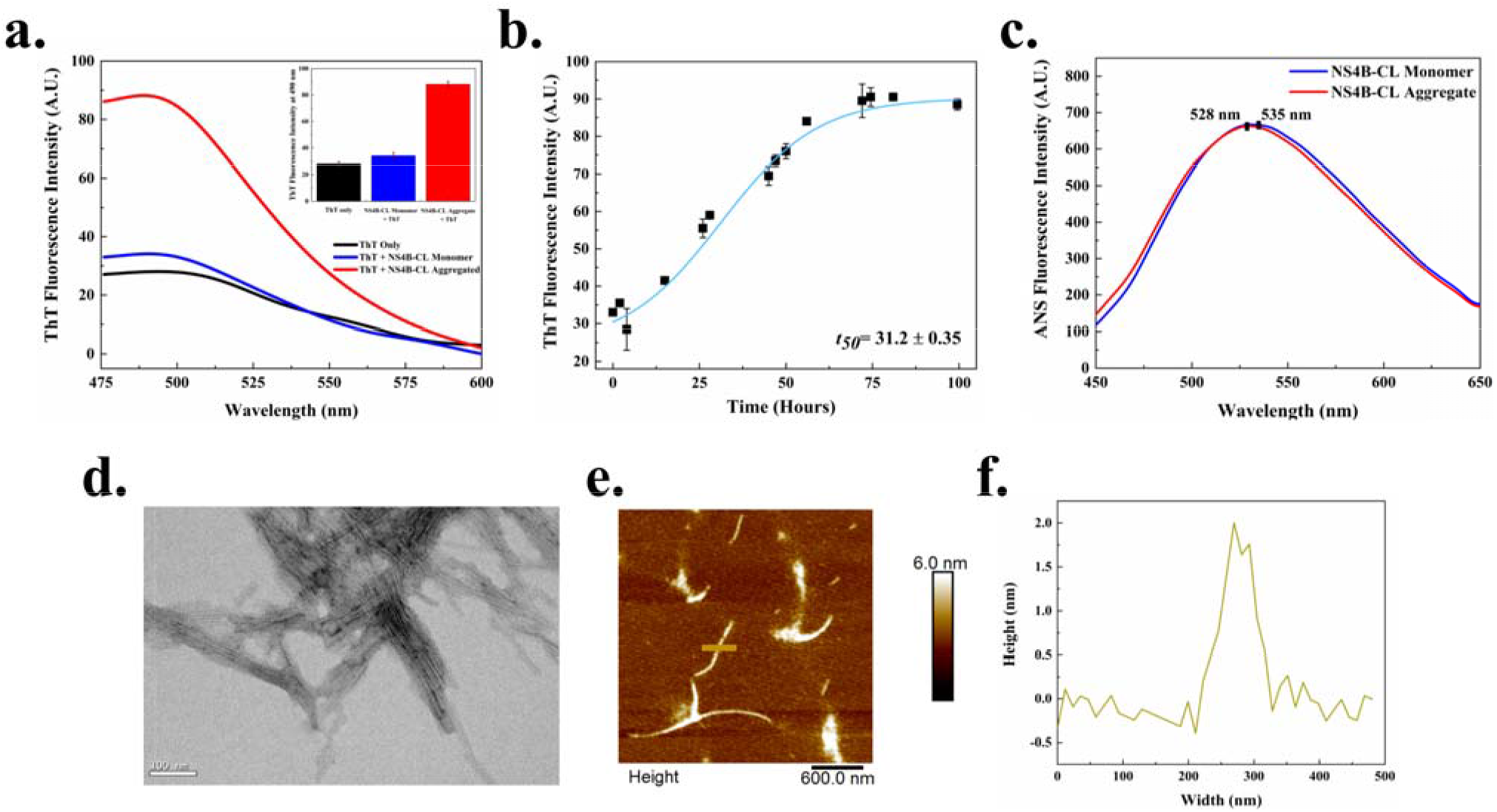
ZIKV NS4B-CR forms amyloid-like fibrils in-vitro. **(a)** ThT fluorescence scan shows a ~three-fold increase in ThT fluorescence at 490 nm after 120 hrs signifies the formation of aggregates. **(b)** represents the aggregation kinetics of NS4B-CR amyloidogenesis where the half time (*t_50_*) is calculated to be 31.2±0.35 hrs. **(c)** Fluorescence maxima of ANS dye also shows a noticeable blue-shift from 535 nm (monomer) to 528 nm (120 hrs aggregate). **(d)** HR-TEM image of 96 hrs aggregates reveals several overlapping and single fibrils. Scale bar represents 100 nm. **(e)** AFM image of 96 hrs aggregates depicts a 3D amyloid fibril having a height of ~2 nm shown in section **(f)**.

## Discussion

The occurrence of aggregation “hot spots” has been well-defined in the proteins and peptides, leading to several neurological diseases. There are more than 50 diseases known so far where protein aggregation and amyloid formation has been implicated as the hallmark.^37,38^ For example, Aβ42, and α-synuclein plaques are the hallmark features of two major systemic amyloidogenic disorders - Alzheimer’s and Parkinson’s diseases, respectively.^39^ Previously, these protein misfolding diseases are found to be affected by the presence of several viruses. HSV-1 and −2, varicella zoster-virus, Ljuangan virus, H5N1 influenza virus, and HIV have been correlated with such acute neurological diseases.^40–45^ HSV-1 infection is found to up-regulate certain mediators of AD as well as cause gliosis and inflammation similar to observed in the disease.^43^ HIV has been demonstrated to increase the levels of Aβ42 in cerebrospinal fluid in HIV-associated neurologic disease, known as HAND.^41^ Further, the Ljuangan virus has been found to be present in neurons, astrocytes, and amyloid plaques in the hippocampus region of Alzheimer’s patients.^40^ A closely-related Hepatitis C virus in extreme cases can cause cognitive impairment and dementia.^46^ Moreover, aggregation of viral proteins has been demonstrated in several cases. PB1-F2 protein of influenza itself adopts β-sheets conformation and forms amorphous aggregates in vitro.^47^ Similarly, the M45 protein of murine cytomegalovirus derives self-assembly into amyloid fibrils. Furthermore, it interacts with many host proteins such as RIPK1 and RIPK3 and forms hetero-oligomeric amyloid fibrils altering their natural functions.^48^ Recently, many protein regions of SARS-CoV-2 are observed to form amyloid-like structures in vitro.^49^ An important thing to note here is that viruses can induce severe neurological symptoms by regulating the pathways vital for neural function and directly or indirectly affect the major hallmark proteins like Aβ42, APP, and Tau. They can also exert their pathogenesis through an aggregation-mediated pathway where the viral proteins are not only able to form insoluble aggregates but also forms hetero-oligomeric fibrils with host proteins. It is a well-known fact that the deposition of insoluble amyloid-like aggregates in between the neural cells can cause brain tissue atrophy. Evidence show that viruses have been evolving to be more pathogenic and are adopting misfolded conformations (viral proteins) while also affecting the host protein-folding pathways. It can be one of the ongoing evolutionary ways which require thorough investigation.

As a neurotropic arbovirus, ZIKV has been associated with many severe neurological disorders of both central and peripheral nervous systems.^50^ Its remarkable ability to invade the placental-blood-barrier, which only a few viruses are capable of, can cause irreversible damage to the fetal brain. Congenital ZIKV syndrome consists of a group of birth defects that includes microcephaly, partly collapsed skull, and hydrocephaly like disorders.^51^ In infants, it persists even after birth which can trigger long-lasting effects on the nervous system and thus might be related to several neurological diseases.^51^ As seen with various distantly related viruses, reports of ZIKV affecting the acute neurological disorders in comorbid conditions have surfaced in the last few years.^52,53^ A study reported the altered expression of neurological disorders-related proteins in human mesenchymal cells by ZIKV infection, providing a possible link between viral infection with AD, Parkinson’s disease, Schizophrenia, and Amyotrophic Lateral Sclerosis.^52^ Further link between AD with ZIKV infection is established by the virus-mediated increased expression of the amyloid precursor protein, a key membrane protein involved in the pathogenesis of AD.^53^

Links between neurological disorders and ZIKV infection in comorbid conditions compelled us to investigate the novel molecular mechanisms employed by the virus for its pathogenicity. Here, we have identified numerous amyloidogenic regions in the ZIKV proteome and have analyzed the aggregation potential of its proteins and protein regions in vitro. Its envelope domain III, NS1 *β*-roll peptide, 2K peptide, and NS4B-CR are demonstrated to form homo-oligomeric amyloid-like fibrillar aggregates. These results show the amyloidogenic potential of ZIKV proteins which might be a way of viral pathogenesis, especially in comorbid conditions.

## Conclusions

The protein misfolding leads to amyloid fibrils which are found to be linked with a number of human diseases. The abnormal deposition of proteins leads to amyloid plaque, causing tissue damage and organ dysfunction. The neurodegenerative disease such as microcephaly linked with ZIKV is still a mystery in terms of its pathological mechanism. The aggregation propensity of ZIKV proteome lays some interesting questions for its pathological functionality, which are hidden till now. The experimental validation of proteins and peptides indicates the high potential of APRs in the ZIKV proteome to form amyloid fibrils. Further, studies are needed to establish a mechanistic link between ZIKV and its associated neurodegenerative pathogenesis. We believe that this present study will have implications in the discovery of some novel molecules against ZIKV originated amyloid fibrils and associated diseases.

## Materials & Methods

### Peptides and chemicals

The peptides corresponding to NS1 *β*-roll, 2K region, NS4B-CR were chemically synthesized and purchased from Thermo Scientific, USA. Other chemicals, including thioflavin T (ThT), 8-Anilinonaphthalene-1-sulfonic acid (ANS), and ammonium molybdate, were purchased from Sigma Aldrich, St. Louis, USA. MICA sheets and TEM Grids were acquired from TED PELLA INC., USA.

### In-silico analysis of aggregation propensity profile

The ZIKV proteome sequences for the strain Mr766 (UniProt ID: Q32ZE1) were derived from the UniProt database. Aggregation propensity prediction was performed by four different tools, namely TANGO^54^, AGGRESCAN^55^, MetAmyl^56^ and FoldAmyloid^57^. For predicting the hydrophilicity of protein CamSol^58^ server was employed. Briefly, in TANGO, the parameters used were that of physiological condition (pH 7.4; Temperature 37 °C) except for the ionic strength, which was left to default. In CamSol, AGGRESCAN, MetAmyl, and FoldAmyloid, complete default parameters were used.

### Prediction of 20S Proteasome Cleavage Sites in ZIKV proteome

ZIKV structural and non-structural proteins were fed into the proteasome cleavage prediction tools Pcleavage and NetChop with threshold/cutoff values of 0.6 and 0.9, respectively.^32,33^

### Cloning and expression of EDIII domain of ZIKV E protein

The protein construct used for experimental studies was “MGVSYSLCTAAFTFTKIPAETLHGTVTVEVQYAGTDGPCKVPAQMAVDMQTLTPVGRLITANPVITESTENSKMMLELDPPFGDSYIVIGVGEKKITHHWHRSGSTIGKGQFYLNELE HHHHHH”. Corresponding to residues 302 – 409 of ZIKV Envelope protein, EDIII was cloned into pET-21(d) expression vector, which was transformed into BL21 (DE3) strain of *E. coli*. Cells were grown in 1 L of LB media up to OD_600_ of 0.5 and EDIII was induced using 1 mM IPTG at 37 °C for 4 hrs. It was purified using Ni-NTA affinity chromatography by the following protocol. The cell pellet was resuspended in resuspension buffer (50 mM Tris-Cl, 300 mM NaCl, 40 mM Imidazole, 5% Glycerol, pH 8) and protease inhibitor cocktail. After sonication, the cell lysate was centrifuged at 13,000 rpm for 30 min, and subsequent pellet was resuspended and washed twice using a washing buffer (25 mM Tris-Cl, 2 M Urea, 1% Triton X-100, pH 8). After centrifugation, the soft pellet was resuspended in an extraction buffer (25 mM Tris-Cl, 300 mM NaCl, 8 M Urea, pH 8) and centrifuged at 13,000 rpm for 60 min. The supernatant containing EDIII was refolded in resuspension buffer, bound to His-tag column, and eluted using an elution buffer (50 mM Tris-Cl, 300 mM NaCl, 1 M Imidazole, 5% Glycerol, pH 8). It was stored in a buffer (20 mM Tris-Cl, 150 mM NaCl, 5% Glycerol, pH 8) in aliquots. The identity of the eluted protein was confirmed using SDS-PAGE and CD spectroscopy.

### Preparation of aggregates

To dissolve any pre-formed aggregates, peptides were treated with HFIP solution (100%). It was left to evaporate at room temperature in a desiccator for overnight. The 2K peptide was dissolved in 10 % DMSO and 20 mM sodium phosphate buffer (pH 7.4) at a 1 mg/ml concentration. The NS1 *β*-roll peptide was dissolved in 1 X PBS (pH 7) at 0.5 mg/ml concentration. The NS4B-CR peptide was dissolved in 20 mM sodium phosphate buffer (pH 7.4) at a 1 mg/ml concentration. The EDIII protein was purified as described above. The peptides and EDIII protein were then subjected to constant shaking at 1000 rpm and 37 °C using an Eppendorf ThermoMixer^®^ C. Samples were collected at different time intervals during experiments.

### Thioflavin T (ThT) fluorescence and kinetics

ThT fluorescent dye binds selectively with amyloid fibrils to give a fluorescence peak at 490 nm.^59,60^ Therefore, it was employed to study the formation of aggregates and kinetics reaction of ZIKV protein regions. Samples were prepared by below given protocol:

| **ZIKV protein and peptides** | **Concentration of ThT dye** | **Concentration of peptides** | **Buffer** | **Control** |
| --- | --- | --- | --- | --- |
| **EDIII** | 10 μM | 10 μM | 50mM Tris-CL<br>(pH 8) | 50mM Tris-CL<br>(pH 8) + 10 μM<br>ThT dye |
| **NS1 β-roll** | 10 μM | 10 μM | 1 X PBS (pH<br>7) | 1 X PBS (pH 7)<br>+ 10 μM ThT<br>dye |
| **2K** | 10 μM | 30 μM | 10 % DMSO in<br>20 mM sodium<br>phosphate (pH<br>7.4) | 10 % DMSO in<br>20 mM sodium<br>phosphate (pH<br>7.4) + 10 μM<br>dye |
| **NS4B-CR** | 20 μM | 15 μM | 20 mM sodium<br>phosphate (pH<br>7.4) | 20mM sodium<br>phosphate (pH<br>7.4) + 20 μM<br>ThT dye |

Samples were incubated for 10 min, after which ThT fluorescence was monitored from 450 nm – 650 nm range on excitation at 440 nm in a flat black 96-well plate using TECAN Infinite^®^ 200 PRO multimode microplate reader. All the measurements were set up in duplicates, and the average value was reported with standard deviation (SD). Furthermore, the kinetics of fibril formation for peptides NS1 *β*-roll and NS4B-CR was studied by plotting the change in ThT fluorescence intensity on the y-axis with time (hrs) on the x-axis. The resultant data plot was fitted using a sigmoidal curve to obtain *t*_*50*_ (when ThT fluorescence intensity reaches 50% of its maximum value). The equation describing the kinetics of fibrillogenesis is:

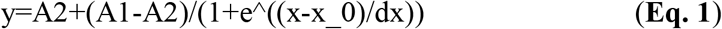

Where A1 indicates the initial fluorescence, A2 is the final fluorescence, x_0 is the midpoint (*t*_*50*_ value), and dx is a time constant.

### 8-Anilinonaphthalene-1-sulfonic acid (ANS) fluorescence

ANS, an extensively utilized fluorescent molecular probe, binds to cationic groups of proteins and is routinely used to study solvent-exposed hydrophobic clusters of proteins during amyloidogenesis.^60^ ZIKV protein regions were monitored for their susceptibility of binding with ANS dye during the aggregation process. Samples were prepared by below given protocol:

| **ZIKV protein and peptides** | **Concentration of ANS dye** | **Concentration of peptides** | **Buffer** | **Control** |
| --- | --- | --- | --- | --- |
| **EDIII** | 30 μM | 10 μM | 50mM Tris-CL (pH 8) | 50mM Tris-CL (pH 8) + 30 μM ANS dye |
| **NS1 β-roll** | 10 μM | 50 μM | 1 X PBS (pH 7) | 1 X PBS (pH 7) + 10 μM ANS dye |
| **2K** | 10 μM | 30 μM | 10 % DMSO and 20 mM sodium phosphate (pH 7.4) | 10 % DMSO in 20 mM sodium phosphate (pH 7.4) + 10 ANS dye |
| **NS4B-CR** | 20 μM | 10 μM | 20 mM sodium phosphate (pH 7.4) | 20mM sodium phosphate (pH 7.4) + 20 μM ANS dye |

Sample mix were incubated for 10 minutes, and fluorescence intensities were documented from 400 nm to 700 nm range on excitation at 380 nm in 1 ml path length quartz cuvette using Horiba Scientific Fluorolog-3.

### Atomic force microscopy (AFM)

AFM images were captured using tapping-mode AFM (Dimension Icon from Bruker) system. Aggregated samples after thirty-fold dilution were drop-casted on the mica surface. The mica sheets were dried at room temperature overnight, and images were captured.

### High-resolution transmission electron microscopy (HR-TEM)

Morphology of aggregates was also confirmed through HR-TEM (FP 5022/22-Tecnai G2 20 S-TWIN, FEI). A twenty-fold diluted solution of aggregated samples were mounted on 200-mesh carbon-coated copper grids (Ted Pella, Inc, USA). Samples were then negatively stained using 3% ammonium molybdate solution, following which grids were air-dried overnight, and images were captured using HR-TEM with an accelerating voltage of 200 kV.

## Data availability

All data are contained within the manuscript or as supporting information.

## Acknowledgment

Authors are thankful to the Indian Institute of Technology Mandi (BioX, AMRC, and C4DFED center) for all the facilities and faculty research grant, School of Basic Sciences, IIT Mandi to RG. KUS and SKK were supported by the Indian Council of Medical Research (ICMR), India, for fellowship. TB is thankful to the Department of Science and Technology (DST) for the INSPIRE Fellowship. RG is grateful for the IYBA Award (Grant Number: BT/11/IYBA/2018/06) from the Department of Biotechnology (DBT), India; MHRD-SPARC (SPARC/2018-2019/P37/SL), Science and Engineering Research Board (SERB), India (Grant Number: CRG/2019/005603); and Indian Council of Medical Research (Grant numbers: 58/6/2020/PHA/BMS, 52/04/2020/BIO/BMS).

## Author contributions

RG: Conception, design, review, data analysis, and writing of the manuscript. TB, KUS, KG, SKK, NS performed experiments and analyzed the data. KUS performed the computational predictions. All authors have contributed to manuscript writing.

## References

1. Musso, D. & Gubler, D. J. Zika Virus. Clinical Microbiology Reviews 29, 487 (2016).

2. G, R., S, Y. & R, K. Zika virus: An overview. Journal of family medicine and primary care 5, 523 (2016).

3. Olmo, I. G. et al. Zika virus promotes neuronal cell death in a non-cell autonomous manner by triggering the release of neurotoxic factors. Frontiers in Immunology 8, 1016 (2017).

4. Yang, S. et al. Zika Virus-Induced Neuronal Apoptosis via Increased Mitochondrial Fragmentation. Frontiers in Microbiology 11, 3316 (2020).

5. Huang, W. C., Abraham, R., Shim, B. S., Choe, H. & Page, D. T. Zika virus infection during the period of maximal brain growth causes microcephaly and corticospinal neuron apoptosis in wild type mice. Scientific Reports 2016 6:1 6, 34793 (2016).

6. CM, D. Protein aggregation and its consequences for human disease. Protein and peptide letters 13, 219–227 (2006).

7. AJ, W., S, I., MJ, T. & CM, G. Distinct stress conditions result in aggregation of proteins with similar properties. Scientific reports 6, (2016).

8. Moshe, A. & Gorovits, R. Virus-Induced Aggregates in Infected Cells. Viruses 4, 2232 (2012).

9. Olasunkanmi, O. I., Chen, S., Mageto, J. & Zhong, Z. Virus-Induced Cytoplasmic Aggregates and Inclusions are Critical Cellular Regulatory and Antiviral Factors. Viruses 12, 399 (2020).

10. Saumya, K. U., Gadhave, K., Kumar, A. & Giri, R. Zika virus capsid anchor forms cytotoxic amyloid-like fibrils. Virology 560, 8–16 (2021).

11. Ashraf, G. M. et al. Protein misfolding and aggregation in Alzheimer’s disease and Type 2 Diabetes Mellitus. CNS & neurological disorders drug targets 13, 1280 (2014).

12. J, G. et al. Protein misfolding and aggregation in neurodegenerative diseases: a review of pathogeneses, novel detection strategies, and potential therapeutics. Reviews in the neurosciences 30, 339–358 (2019).

13. Jucker, M. & Walker, L. C. Propagation and spread of pathogenic protein assemblies in neurodegenerative diseases. Nature neuroscience 21, 1341 (2018).

14. J, S., J, P., I, P., V, I. & S, V. Computational prediction of protein aggregation: Advances in proteomics, conformation-specific algorithms and biotechnological applications. Computational and structural biotechnology journal 18, 1403–1413 (2020).

15. TY, T. et al. Capsid protein structure in Zika virus reveals the flavivirus assembly process. Nature communications 11, (2020).

16. KU, S., K, G., A, K. & R, G. Zika virus capsid anchor forms cytotoxic amyloid-like fibrils. Virology 560, 8–16 (2021).

17. R, B., MK, C. & ML, N. Specific interaction of capsid protein and importin-alpha/beta influences West Nile virus production. Biochemical and biophysical research communications 389, 63–69 (2009).

18. J, R. et al. Role of Capsid Anchor in the Morphogenesis of Zika Virus. Journal of virology 92, (2018).

19. L, D. et al. Structures of the Zika Virus Envelope Protein and Its Complex with a Flavivirus Broadly Protective Antibody. Cell host & microbe 19, 696–704 (2016).

20. Nambala, P. & Su, W.-C. Role of Zika Virus prM Protein in Viral Pathogenicity and Use in Vaccine Development. Frontiers in Microbiology 9, 1797 (2018).

21. Zhang, X., Jia, R., Shen, H., Wang, M. & Yin, Z. Structures and Functions of the Envelope Glycoprotein in Flavivirus Infections. 1–14 (2017). doi:10.3390/v9110338

22. A, W., S, T., L, I., K, H. & R, H. Zika virus genome biology and molecular pathogenesis. Emerging microbes & infections 6, (2017).

23. M, R., N, S. & SK, S. Flavivirus NS1: a multifaceted enigmatic viral protein. Virology journal 13, (2016).

24. DL, A., WC, B., J, J., RJ, K. & JL, S. Structure-guided insights on the role of NS1 in flavivirus infection. BioEssays□: news and reviews in molecular, cellular and developmental biology 37, 489–494 (2015).

25. JY, L. et al. Role of nonstructural protein NS2A in flavivirus assembly. Journal of virology 82, 4731–4741 (2008).

26. J, L. et al. Crystal structure of Zika virus NS2B-NS3 protease in complex with a boronate inhibitor. Science (New York, N.Y.) 353, 503–505 (2016).

27. R, A. et al. Crystal structure of a novel conformational state of the flavivirus NS3 protein: implications for polyprotein processing and viral replication. Journal of virology 83, 12895–12906 (2009).

28. H, N., F, P.-J. & J, V. NS4A and NS4B proteins from dengue virus: membranotropic regions. Biochimica et biophysica acta 1818, 2818–2830 (2012).

29. J, Z. et al. Characterization of dengue virus NS4A and NS4B protein interaction. Journal of virology 89, 3455–3470 (2015).

30. D, K., A, K., T, B. & R, G. Zika virus NS4A N-Terminal region (1-48) acts as a cofactor for inducing NTPase activity of NS3 helicase but not NS3 protease. Archives of biochemistry and biophysics 695, (2020).

31. AK, U. et al. Crystal structure of full-length Zika virus NS5 protein reveals a conformation similar to Japanese encephalitis virus NS5. Acta crystallographica. Section F, Structural biology communications 73, 116–122 (2017).

32. M, B. & GP, R. Pcleavage: an SVM based method for prediction of constitutive proteasome and immunoproteasome cleavage sites in antigenic sequences. Nucleic acids research 33, (2005).

33. C, K., AK, N., H, S., V, D. & S, B. Prediction of proteasome cleavage motifs by neural networks. Protein engineering 15, 287–296 (2002).

34. Tai, W. et al. Rational Design of Zika Virus Subunit Vaccine with Enhanced Efficacy. Journal of Virology 93, (2019).

35. Zou, G. et al. A single-amino acid substitution in West Nile virus 2K peptide between NS4A and NS4B confers resistance to lycorine, a flavivirus inhibitor. Virology 384, 242–252 (2009).

36. J, Z., J, N. & K, D. Flaviviral NS4b, chameleon and jack-in-the-box roles in viral replication and pathogenesis, and a molecular target for antiviral intervention. Reviews in medical virology 25, 205–223 (2015).

37. Skretas, G. & Ventura, S. Editorial: Protein Aggregation and Solubility in Microorganisms (Archaea, Bacteria and Unicellular Eukaryotes): Implications and Applications. Frontiers in Microbiology 620239 (2020). doi:10.3389/FMICB.2020.620239/PDF

38. Chiti, F. & Dobson, C. M. Protein Misfolding, Amyloid Formation, and Human Disease: A Summary of Progress Over the Last Decade. Annual review of biochemistry 86, 27–68 (2017).

39. Groot, N. S. de, Pallarés, I., Avilés, F. X., Vendrell, J. & Ventura, S. Prediction of ‘hot spots’ of aggregation in disease-linked polypeptides. BMC Structural Biology 5, 18 (2005).

40. Niklasson, B., Lindquist, L., Klitz, W. & Englund, E. Picornavirus Identified in Alzheimer’s Disease Brains: A Pathogenic Path? Journal of Alzheimer’s Disease Reports 4, 141–146 (2020).

41. Clifford, D. B. et al. CSF biomarkers of Alzheimer disease in HIV-associated neurologic disease. Neurology 73, 1982–1987 (2009).

42. Nimgaonkar, V. L. et al. Temporal cognitive decline associated with exposure to infectious agents in a population-based, aging cohort. Alzheimer Disease and Associated Disorders 30, 216–222 (2016).

43. Cairns, D. M. et al. A 3D human brain–like tissue model of herpes-induced Alzheimer’s disease. Science Advances 6, eaay8828 (2020).

44. Jang, H. et al. Highly pathogenic H5N1 influenza virus can enter the central nervous system and induce neuroinflammation and neurodegeneration. Proceedings of the National Academy of Sciences of the United States of America 106, 14063–14068 (2009).

45. De Clercq, E. & Li, G. Approved antiviral drugs over the past 50 years. Clinical Microbiology Reviews 29, 695–747 (2016).

46. WC, C. et al. Hepatitis C viral infection and the risk of dementia. European journal of neurology 21, 1068–e59 (2014).

47. Chevalier, C. et al. PB1-F2 influenza A virus protein adopts a β-sheet conformation and forms amyloid fibers in membrane environments. Journal of Biological Chemistry 285, 13233–13243 (2010).

48. Pham, C. L. et al. Viral M45 and necroptosis-associated proteins form heteromeric amyloid assemblies. EMBO reports 20, (2019).

49. Bhardwaj, T. et al. Amyloidogenic proteins in the SARS-CoV and SARS-CoV-2 proteomes. bioRxiv 2021.05.29.446267 (2021). doi:10.1101/2021.05.29.446267

50. Blázquez, A.-B. & Saiz, J.-C. Neurological manifestations of Zika virus infection. World Journal of Virology 5, 135 (2016).

51. Wheeler, A. C. Development of Infants With Congenital Zika Syndrome: What Do We Know and What Can We Expect? Pediatrics 141, S154 (2018).

52. WO, B.-S. et al. Zika Virus Infection of Human Mesenchymal Stem Cells Promotes Differential Expression of Proteins Linked to Several Neurological Diseases. Molecular neurobiology 56, 4708–4717 (2019).

53. A, L. et al. Amyloid precursor protein is a restriction factor that protects against Zika virus infection in mammalian brains. The Journal of biological chemistry 295, 17114–17127 (2020).

54. Fernandez-Escamilla, A. M., Rousseau, F., Schymkowitz, J. & Serrano, L. Prediction of sequence-dependent and mutational effects on the aggregation of peptides and proteins. Nature Biotechnology 22, 1302–1306 (2004).

55. Conchillo-Solé, O. et al. AGGRESCAN: A server for the prediction and evaluation of ‘hot spots’ of aggregation in polypeptides. BMC Bioinformatics 8, (2007).

56. Emily, M., Talvas, A. & Delamarche, C. MetAmyl: A METa-predictor for AMYLoid proteins. PLoS ONE 8, (2013).

57. Garbuzynskiy, S. O., Lobanov, M. Y. & Galzitskaya, O. V. FoldAmyloid: A method of prediction of amyloidogenic regions from protein sequence. Bioinformatics 26, 326–332 (2009).

58. P, S. & M, V. Protein Solubility Predictions Using the CamSol Method in the Study of Protein Homeostasis. Cold Spring Harbor perspectives in biology 11, (2019).

59. C, X., TY, L., D, C. & Z, G. Thioflavin T as an amyloid dye: fibril quantification, optimal concentration and effect on aggregation. Royal Society open science 4, (2017).

60. Sulatsky, M. I. et al. Effect of the fluorescent probes ThT and ANS on the mature amyloid fibrils. Prion 14, 67–75 (2020).

