## Supplementary File for "The aggregation potential of Zika virus proteome"

**Running title:** Aggregation prone regions in Zika virus

**Table S1: *In-silico* mapping of 20S proteasome cleavage sites across ZIKV proteome predicted with SVM-based method Pcleavage and proteasome prediction server NetChop.** The cleavage site residues are colored in red, and the aggregation-prone regions predicted by TANGO are depicted as blue italicized, underlined sequences.

| **Protein Name,**  **Prediction**  **method** | **Distribution of predicted 20s proteasome cleavage sites** | **Distribution of cleavage sites and Aggregation prone regions (APR)** |
| --- | --- | --- |
| **Structural Proteins** | | |
| Capsid  (C)  (Pcleavage) | 1-MKNPKEEIRRIRIVNMLKRGVARVNPLGGL-30  31-KRLPAGLLLGHGPIR*MVLAILAFL*RFTAIK-60  61-PSLGLINRWGSVGKKEAMEIIKKFKKDLAA-90  91-MLRIINARKERKRR-104 | Total cleavage sites= 2  Sites inside APR= 2 (100%)  Sites outside APR= 0 (0%) |
| Capsid  (C)  (NetChop) | 1-MKNPKEEIRRIRIVNMLKRGVARVNPLGGL-30  31-KRLPAGLLLGHGPIR*MVLAILAFL*RFTAIK-60  61-PSLGLINRWGSVGKKEAMEIIKKFKKDLAA-90  91-MLRIINARKERKRR-104 | Total cleavage sites= 17  Sites inside APR= 4 (23.5%)  Sites outside APR= 13 (76.5%) |
| Capsid Anchor  (CA),  (Pcleavage) | 1-GADTS*IGIIGLLLTTAM*A-18 | Total cleavage sites= 1  Sites inside APR= 1 (100%)  Sites outside APR= 0 (0%) |
| Capsid Anchor  (CA),  NetChop | 1-GADTS*IGIIGLLLTTAM*A-18 | Total cleavage sites= 2  Sites inside APR= 2 (100%)  Sites outside APR= 0 (0%) |
| PrM  (Pcleavage) | 1-AEITRRGS*AYYMYL*DRSDAGKAISFATTLG-30  31-VNKCHVQIMDLGHMCDATMSYECPMLDEGV-60  61-EPDDVDCWCNT*TSTWVVYGT*CHHKKGEARR-90  91-SRRAVTLPSHSTRKLQTRSQTWLESREYTK-120  121-HLIKVENWIFRNP*GFALVAVAIAWLLG*SST-150  151-SQK*VIYLVMILLI*APAYS-168 | Total cleavage sites= 9  Sites inside APR= (55.5%)  Sites outside APR= (44.5%) |
| PrM  (NetChop) | 1-AEITRRGS*AYYMYL*DRSDAGKAISFATTLG-30  31-VNKCHVQIMDLGHMCDATMSYECPMLDEGV-60  61-EPDDVDCWCNT*TSTWVVYGT*CHHKKGEARR-90  91-SRRAVTLPSHSTRKLQTRSQTWLESREYTK-120  121-HLIKVENWIFRNP*GFALVAVAIAWLLG*SST-150  151-SQK*VIYLVMILLI*APAYS-168 | Total cleavage sites= 33  Sites inside APR= 13 (39.4%)  Sites outside APR= 20 (60.6%) |
| Envelope  (E),  (Pcleavage) | 1-IRCIGVSNRDFVEGMSGGTWVDVVLEHGGC-30  31-VTVMAQDKPTVDIELVTTTVSNMAEVRSYC-60  61-YEASISDMASDSRCPTQGEAYLDKQSDTQY-90  91-VCKRTLVDRGWGNGCGLFGKGSLVTCAKFT-120  121-CSKKMTGKSIQPENLEYRIMLSVHGSQHSG-150  151-MIGYETDEDRAKVEVTPNSPRAEATLGGFG-180  181-SLGLDCEPRTGLDFSD*LYYLT*MNNKHWLVH-210  211-KEWFHDIPLPWHAGADTGTPHWNNKEALVE-240  241-FKDAHAKRQ*TVVVLG*SQEGAVHTALAGALE-270  271-AEMDGAKGRLFSGHLKCRLKMDKLRLKGVS-300  301-YSLCTAAFTFTKVPAETLHGTVTVEVQYAG-330  331-TDGPCKIPVQMAVDMQTLTPVGRLITANPV-360  361-ITESTENSKMMLELDPPFGD*SYIVIGVG*DK-390  391-KITHHWHRSGSTIGKAFEATVRGAKRMAVL-420  421-GDTAWDFGSVGGVFNSLGKGIHQIFGAAFK-450  451-SLFGGMS*WFSQILIGTLLVWLGL*NTKNGS*I*-480  481-*SLTCLALGGVMIFLSTAV*SA-500 | Total cleavage sites= 17  Sites inside APR= 1 (5.9%)  Sites outside APR= 16 (94.1%) |
| Envelope  (E),  (NetChop) | 1-IRCIGVSNRDFVEGMSGGTWVDVVLEHGGC-30  31-VTVMAQDKPTVDIELVTTTVSNMAEVRSYC-60  61-YEASISDMASDSRCPTQGEAYLDKQSDTQY-90  91-VCKRTLVDRGWGNGCGLFGKGSLVTCAKFT-120  121-CSKKMTGKSIQPENLEYRIMLSVHGSQHSG-150  151-MIGYETDEDRAKVEVTPNSPRAEATLGGFG-180  181-SLGLDCEPRTGLDFSD*LYYLT*MNNKHWLVH-210  211-KEWFHDIPLPWHAGADTGTPHWNNKEALVE-240  241-FKDAHAKRQ*TVVVLG*SQEGAVHTALAGALE-270  271-AEMDGAKGRLFSGHLKCRLKMDKLRLKGVS-300  301-YSLCTAAFTFTKVPAETLHGTVTVEVQYAG-330  331-TDGPCKIPVQMAVDMQTLTPVGRLITANPV-360  361-ITESTENSKMMLELDPPFGD*SYIVIGVG*DK-390  391-KITHHWHRSGSTIGKAFEATVRGAKRMAVL-420  421-GDTAWDFGSVGGVFNSLGKGIHQIFGAAFK-450  451-SLFGGMS*WFSQILIGTLLVWLGL*NTKNGS*I*-480  481-*SLTCLALGGVMIFLSTAV*SA-500 | Total cleavage sites= 60  Sites inside APR= 9 (15%)  Sites outside APR= 51 (85%) |
| **Non-Structural Proteins** | | |
| Non-structural 1  (NS1),  (Pcleavage) | 1-DVGCSVDFSKKETRCG*TGVFIY*NDVEAWRD-30  31-RYKYHPDSPRRLAAAVKQAWEEGICGISSV-60  61-SRMENIMWKSVEGELNAILEENG*VQLTVVV*-90  91-*G*SVKNPMWRGPQRLPVPVNELPHGWKAWGK-120  121-SYFVRAAKTNNSFVVDGDTLKECPLEHRAW-150  151-NSFLVEDHGFGVFHTSVWLKVREDYSLECD-180  181-P*AVIGTAV*KGREAAHSDLGYWIESEKNDTW-210  211-RLKRAHLIEMKTCEWPKSHTLWTDGVEESD-240  241-LIIPKSLAGPLSHHNTREGYRTQVKGPWHS-270  271-EELEIRFEECPGTKVYVEETCGTRGPSLRS-300  301-TTASGRVIEEWCCRECTMPPLSFRAKDGCW-330  331-YGMEIRPRKEPESNLVRSMVTA-352 | Total cleavage sites= 8  Sites inside APR= 1 (12.5%)  Sites outside APR= 7 (87.5%) |
| Non-structural 1  (NS1),  (NetChop) | 1-DVGCSVDFSKKETRCG*TGVFIY*NDVEAWRD-30  31-RYKYHPDSPRRLAAAVKQAWEEGICGISSV-60  61-SRMENIMWKSVEGELNAILEENG*VQLTVVV*-90  91-*G*SVKNPMWRGPQRLPVPVNELPHGWKAWGK-120  121-SYFVRAAKTNNSFVVDGDTLKECPLEHRAW-150  151-NSFLVEDHGFGVFHTSVWLKVREDYSLECD-180  181-P*AVIGTAV*KGREAAHSDLGYWIESEKNDTW-210  211-RLKRAHLIEMKTCEWPKSHTLWTDGVEESD-240  241-LIIPKSLAGPLSHHNTREGYRTQVKGPWHS-270  271-EELEIRFEECPGTKVYVEETCGTRGPSLRS-300  301-TTASGRVIEEWCCRECTMPPLSFRAKDGCW-330  331-YGMEIRPRKEPESNLVRSMVTA-352 | Total cleavage sites= 42  Sites inside APR= 3 (7.1%)  Sites outside APR= 39 (92.9%) |
| Non-structural 2A  (NS2A),  (Pcleavage) | 1-GSTDHMDH*FSLGVLVILLMV*QEGLKKRMTT-30  31-K*IIMSTSMAVLVVMILGGFS*MSDLAK*LVIL*-60  61-*MGATFA*EMNTGGD*VAHLALVAAF*KVRP*ALL*-90  91-*VSFIF*RANWTPRE*SMLLALASCL*LQTAISA-120  121-LEGD*LMVLINGFALAWLAI*RAMAVPRTDNI-150  151-ALP*ILAAL*TPLARGT*LLVAW*RAGLATCGGI-180  181-MLLSLKGKGSVKKNLP*FVMALGLTAV*RVVD-210  211-P*INVVGLLLLT*RSGKR-226 | Total cleavage sites= 12  Sites inside APR= 8 (66.7%)  Sites outside APR= 4 (33.3%) |
| Non-structural 2A  (NS2A),  (NetChop) | 1-GSTDHMDH*FSLGVLVILLMV*QEGLKKRMTT-30  31-K*IIMSTSMAVLVVMILGGFS*MSDLAK*LVIL*-60  61-*MGATFA*EMNTGGD*VAHLALVAAF*KVRP*ALL*-90  91-*VSFIF*RANWTPRE*SMLLALASCL*LQTAISA-120  121-LEGD*LMVLINGFALAWLAI*RAMAVPRTDNI-150  151-ALP*ILAAL*TPLARGT*LLVAW*RAGLATCGGI-180  181-MLLSLKGKGSVKKNLP*FVMALGLTAV*RVVD-210  211-P*INVVGLLLLT*RSGKR-226 | Total cleavage sites= 45  Sites inside APR= 28 (62.2%)  Sites outside APR= 17 (37.8%) |
| Non-structural 2B  (NS2B),  (Pcleavage) | 1-SWPPSE*VLTAVGLICALAGGFA*KADIEMAG-30  31-P*MAAVGLLIVSYVV*SGKSVDMYIERAGDIT-60  61-WEKDAEVTGNSPRLDVALDESGDFSLVEED-90  91-GPPMR*EIILKVVLMAIC*GMNPIAIP*FAAGA*-120  121-*WYVYV*KTGKR-130 | Total cleavage sites= 6  Sites inside APR= 5 (83.3%)  Sites outside APR= 1 (16.7%) |
| Non-structural 2B  (NS2B),  (NetChop) | 1-SWPPSE*VLTAVGLICALAGGFA*KADIEMAG-30  31-P*MAAVGLLIVSYVV*SGKSVDMYIERAGDIT-60  61-WEKDAEVTGNSPRLDVALDESGDFSLVEED-90  91-GPPMR*EIILKVVLMAIC*GMNPIAIP*FAAGA*-120  121-*WYVYV*KTGKR-130 | Total cleavage sites= 18  Sites inside APR= 11 (61.2%)  Sites outside APR= 7 (38.8%) |
| Non-structural NS3  (NS3),  (Pcleavage) | 1-SGALWDVPAPKEVKKGETTDGVYRVMTRRL-30  31-LGSTQVGVGVMQEGVFHTMWHVTKGAALRS-60  61-GEGRLDPYWGDVKQDLVSYCGPWKLDAAWD-90  91-GLSEVQLLAVPPGERARNIQTLPGIFKTKD-120  121-GDIGAVALDYPAGTSGSPILDKCGRVIGLY-150  151-GNGVVIKNGSYVSAITQGKREEETPVECFE-180  181-PSMLKKKQLTVLDLHPGAGKTRRVLPEIVR-210  211-EAIKKRLRTVILAPTRVVAAEMEEALRGLP-240  241-VRYMTTAVNVTHSGTEIVDLMCHATFTSRL-270  271-LQPIRVPNYNLNIMDEAHFTDPSSIAARGY-300  301-ISTRVEMGE*AAAIFMTA*TPPGTRDAFPDSN-330  331-SPIMDTEVEVPERAWSSGFDWVTDHSGKTV-360  361-WFVPSVRNGNEIAACLTKAGKRVIQLSRKT-390  391-FETEFQKTKNQEWDFVITTDISEMGANFKA-420  421-DRVIDSRRCLKPVILDGERVILAGPMPVTH-450  451-ASAAQRRGRIGRNPNKPGDEYMYGGGCAET-480  481-DEGHAHWLEARMLLDNIYLQDG*LIASLY*RP-510  511-EADKVAAIEGEFKLRTEQRKTFVELMKRGD-540  541-LP*VWLAYQVASAGITYT*DRRWCFDGTTNNT-570  571-IMEDSVPAEVWTKYGEKRVLKPRWMDARVC-600  601-SDHAALKSFKEFAAGKR-617 | Total cleavage sites= 20  Sites inside APR= 5 (25%)  Sites outside APR=20 (75%) |
| Non-structural NS3  (NS3),  (NetChop) | 1-SGALWDVPAPKEVKKGETTDGVYRVMTRRL-30  31-LGSTQVGVGVMQEGVFHTMWHVTKGAALRS-60  61-GEGRLDPYWGDVKQDLVSYCGPWKLDAAWD-90  91-GLSEVQLLAVPPGERARNIQTLPGIFKTKD-120  121-GDIGAVALDYPAGTSGSPILDKCGRVIGLY-150  151-GNGVVIKNGSYVSAITQGKREEETPVECFE-180  181-PSMLKKKQLTVLDLHPGAGKTRRVLPEIVR-210  211-EAIKKRLRTVILAPTRVVAAEMEEALRGLP-240  241-VRYMTTAVNVTHSGTEIVDLMCHATFTSRL-270  271-LQPIRVPNYNLNIMDEAHFTDPSSIAARGY-300  301-ISTRVEMGE*AAAIFMTA*TPPGTRDAFPDSN-330  331-SPIMDTEVEVPERAWSSGFDWVTDHSGKTV-360  361-WFVPSVRNGNEIAACLTKAGKRVIQLSRKT-390  391-FETEFQKTKNQEWDFVITTDISEMGANFKA-420  421-DRVIDSRRCLKPVILDGERVILAGPMPVTH-450  451-ASAAQRRGRIGRNPNKPGDEYMYGGGCAET-480  481-DEGHAHWLEARMLLDNIYLQDG*LIASLY*RP-510  511-EADKVAAIEGEFKLRTEQRKTFVELMKRGD-540  541-LP*VWLAYQVASAGITYT*DRRWCFDGTTNNT-570  571-IMEDSVPAEVWTKYGEKRVLKPRWMDARVC-600  601-SDHAALKSFKEFAAGKR-617 | Protein length= 617 aa.  Total cleavage sites= 85  Sites inside APR= 5 (5.9%)  Sites outside APR= 80 (94.1%) |
| Non-structural 4A  (NS4A),  (Pcleavage) | 1-GAALGVMEALGTLPGHMTERFQEAIDN*LAV*-30  31-*LM*RAETGSRPYKAAAAQLPETLET*IMLLGL*-60  61-*LGTVSLGIFFVL*MRNKGIGKMG*FGMVTLGA*-90  91-*SAWLMWL*SEIEPAR*IACVLIVVFLLLVVLI-*120  121-*P*EPEKQR-127 |  |
| Non-structural 4A  (NS4A),  (NetChop) | 1-GAALGVMEALGTLPGHMTERFQEAIDN*LAV*-30  31-*LM*RAETGSRPYKAAAAQLPETLET*IMLLGL*-60  61-*LGTVSLGIFFVL*MRNKGIGKMG*FGMVTLGA*-90  91-*SAWLMWL*SEIEPAR*IACVLIVVFLLLVVLI-*120  121-*P*EPEKQR-127 | Protein length= 127 aa.  Total cleavage sites= 22  Sites inside APR= 16 (72.7%)  Sites outside APR= 6 (27.3%) |
| 2K,  (Pcleavage) | 1-SPQDNQ*MAIIIMVAVGLLGLIT*A-23 |  |
| 2K,  (NetChop) | 1-SPQDNQ*MAIIIMVAVGLLGLIT*A-23 | Protein length= 23 aa.  Total cleavage sites= 4  Sites inside APR= 4 (100%)  Sites outside APR= 0 (0%) |
| Non-structural 4B  (NS4B),  (Pcleavage) | 1-NELGWLERTKNDIAHLMGRREEGATMGFSM-30  31-DIDLRPA*SAWAIYAALTTLI*TPAVQHAVTT-60  61-SYNNYSLMAMATQAGVLFGMGKGMPFMHGD-90  91-LGVPLLMMGCYSQLTP*LTLIVAIILLVAHY*-120  121-*MYL*IPGLQAAAARAAQKRTAAGIMKNPVVD-150  151-GIVVTDIDTMTIDPQVEKKMGQ*VLLIAVAI*-180  181-*SSAVLL*RTAWGWGEAGALITAATSTLWEGS-210  211-PNKYWNSSTATSLCNIFRGSY*LAGASLIYT*-240  241-*VT*RNAGLVKRR-251 |  |
| Non-structural 4B  (NS4B),  (NetChop) | 1-NELGWLERTKNDIAHLMGRREEGATMGFSM-30  31-DIDLRPA*SAWAIYAALTTLI*TPAVQHAVTT-60  61-SYNNYSLMAMATQAGVLFGMGKGMPFMHGD-90  91-LGVPLLMMGCYSQLTP*LTLIVAIILLVAHY*-120  121-*MYL*IPGLQAAAARAAQKRTAAGIMKNPVVD-150  151-GIVVTDIDTMTIDPQVEKKMGQ*VLLIAVAI*-180  181-*SSAVLL*RTAWGWGEAGALITAATSTLWEGS-210  211-PNKYWNSSTATSLCNIFRGSY*LAGASLIYT*-240  241-*VT*RNAGLVKRR-251 | Protein length= 251 aa.  Total cleavage sites= 44  Sites inside APR= 17 (38.6%)  Sites outside APR= 27 (61.4%) |
| Non-structural 5  (NS5),  (Pcleavage) | 1-GGGTGETLGEKWKARLNQMSALEFYSYKKS-30  31-GITEVCREEARRALKDGVATGGHAVSRGSA-60  61-KIRWLEERGYLQPYGKVVDLGCGRGGWSYY-90  91-AATIRKVQEVRGYTKGGPGHEEPMLVQSYG-120  121-WNIVRLKSGVDVFHMAAEPCDTLLCDIGES-150  151-SSSPEVEETRTLR*VLSMV*GDWLEKRPGAFC-180  181-IKVLCPYTSTMMETMERLQRRHGGGLVRVP-210  211-LCRNSTHEMYWVSGAKSNIIKSVSTTSQLL-240  241-LGRMDGPRRPVKYEEDVNLGSGTRAVASCA-270  271-EAPNMKIIGRRIERIRNEHAETWFLDENHP-300  301-YRTWAYHGSYEAPTQGSASSLVNGVVRLLS-330  331-KPWDVVTGVTGIAMTDTTPYGQQRVFKEKV-360  361-DTRVPDPQEGTRQVMN*IVSSWLW*KELGKRK-390  391-RPRVCTKEEFINKVRSNAALGAIFEEEKEW-420  421-KTAVEAVNDPR*FWALVD*REREHHLRGECHS-450  451-CVYNMMGKREKKQGEFGKAKGSR*AIWYMWL*-480  481-*GARFLEFE*ALGFLNEDHWMGRENSGGGVEG-510  511-LGLQRLGYILEEMNRAPGGKMYADDTAGWD-540  541-TRISKFDLENEALITNQMEEGHR*TLALAVI*-570  571-KYTYQNKVVKVLRPAEGGKTVMDIISRQDQ-600  601-RGSGQ*VVTYALNTFTNLVVQLI*RNMEAEEV-630  631-LEMQDLWLLRKPEKVTRWLQSNGWDRLKRM-660  661-AVSGDDCVVKPIDDRFAHALRFLNDMGKVR-690  691-KDTQEWKPSTGWSNWEEVPFCSHHFNKLYL-720  721-KDGRSIVVPCRHQDELIGRARVSPGAGWSI-750  751-RETACLAKSYAQ*MWQLLYF*HRRDLRLMANA-780  781-ICSAVPVDWVPTGRTTWSIHGKGEWMTTED-810  811-*MLMVW*NRVWIEENDHMEDKTPVTKWTDIPY-840  841-LGKREDLWCGSLIGHRPRTTWAENIKDTVN-870  871-MVRRIIGDEEKYMDYLSTQVRYLGEEGSTP-900  901-GVL-903 |  |
| Non-structural 5  (NS5),  (NetChop) | 1-GGGTGETLGEKWKARLNQMSALEFYSYKKS-30  31-GITEVCREEARRALKDGVATGGHAVSRGSA-60  61-KIRWLEERGYLQPYGKVVDLGCGRGGWSYY-90  91-AATIRKVQEVRGYTKGGPGHEEPMLVQSYG-120  121-WNIVRLKSGVDVFHMAAEPCDTLLCDIGES-150  151-SSSPEVEETRTLR*VLSMV*GDWLEKRPGAFC-180  181-IKVLCPYTSTMMETMERLQRRHGGGLVRVP-210  211-LCRNSTHEMYWVSGAKSNIIKSVSTTSQLL-240  241-LGRMDGPRRPVKYEEDVNLGSGTRAVASCA-270  271-EAPNMKIIGRRIERIRNEHAETWFLDENHP-300  301-YRTWAYHGSYEAPTQGSASSLVNGVVRLLS-330  331-KPWDVVTGVTGIAMTDTTPYGQQRVFKEKV-360  361-DTRVPDPQEGTRQVMN*IVSSWLW*KELGKRK-390  391-RPRVCTKEEFINKVRSNAALGAIFEEEKEW-420  421-KTAVEAVNDPR*FWALVD*REREHHLRGECHS-450  451-CVYNMMGKREKKQGEFGKAKGSR*AIWYMWL*-480  481-*GARFLEFE*ALGFLNEDHWMGRENSGGGVEG-510  511-LGLQRLGYILEEMNRAPGGKMYADDTAGWD-540  541-TRISKFDLENEALITNQMEEGHR*TLALAVI*-570  571-KYTYQNKVVKVLRPAEGGKTVMDIISRQDQ-600  601-RGSGQ*VVTYALNTFTNLVVQLI*RNMEAEEV-630  631-LEMQDLWLLRKPEKVTRWLQSNGWDRLKRM-660  661-AVSGDDCVVKPIDDRFAHALRFLNDMGKVR-690  691-KDTQEWKPSTGWSNWEEVPFCSHHFNKLYL-720  721-KDGRSIVVPCRHQDELIGRARVSPGAGWSI-750  751-RETACLAKSYAQ*MWQLLYF*HRRDLRLMANA-780  781-ICSAVPVDWVPTGRTTWSIHGKGEWMTTED-810  811-*MLMVW*NRVWIEENDHMEDKTPVTKWTDIPY-840  841-LGKREDLWCGSLIGHRPRTTWAENIKDTVN-870  871-MVRRIIGDEEKYMDYLSTQVRYLGEEGSTP-900  901-GVL-903 | Protein length= 903 aa.  Total cleavage sites= 150  Sites inside APR= 27 (18%)  Sites outside APR= 123 (82%) |
